# Biochemical reconstitution defines new functions for membrane-bound glycosidases in assembly of the bacterial cell wall

**DOI:** 10.1101/2021.03.06.434200

**Authors:** Atsushi Taguchi, Suzanne Walker

## Abstract

The peptidoglycan cell wall is a macromolecular structure that encases bacteria and is essential for their survival. Proper assembly of the cell wall requires peptidoglycan synthases as well as membrane-bound cleavage enzymes that control where new peptidoglycan is made and inserted. We are only beginning to understand the roles of peptidoglycan cleavage enzymes in cell wall assembly. Previous studies have shown that two membrane-bound proteins in *Streptococcus pneumoniae*, here named MpgA and MpgB, are important in maintaining cell wall integrity. MpgA was predicted to be a lytic transglycosylase based on its homology to *Escherichia coli* MltG while the enzymatic activity of MpgB was unclear. Using nascent peptidoglycan substrates synthesized *in vitro* from the peptidoglycan precursor Lipid II, we report that both MpgA and MpgB are muramidases. We show that replacing a single amino acid in *E. coli* MltG with the corresponding amino acid from MpgA results in muramidase activity, allowing us to predict from the presence of this amino acid that other putative lytic transglycosylases actually function as muramidases. Strikingly, we report that MpgA and MpgB cut nascent peptidoglycan at different positions along the sugar backbone relative to the reducing end. MpgA produces much longer peptidoglycan oligomers and we show that its cleavage site selectivity is controlled by the LysM-like subdomain, which is also present in MltG. We propose that MltG’s ability to complement loss of MpgA in *S. pneumonia*e despite performing different cleavage chemistry is because it can cleave nascent peptidoglycan at the same distance from the lipid anchor.

## INTRODUCTION

The peptidoglycan cell wall is a major component of the bacterial cell envelope that defines the size and shape of bacteria.^1^ For the bacteria to constantly grow and divide while maintaining proper morphology, the activities of enzymes responsible for peptidoglycan assembly and modification need to be coordinated. Two major families of peptidoglycan synthases have been identified to date. The SEDS (shape, elongation, division and sporulation) proteins are peptidoglycan glycosyltransferases that polymerize the lipid-linked peptidoglycan precursor Lipid II to make glycan strands (Figure 1a).^2,3^ SEDS proteins form a complex with class B penicillin-binding proteins (bPBPs), which are monofunctional transpeptidases that crosslink the glycan strands into the peptidoglycan matrix.^3,4^ The bifunctional class A PBPs (aPBPs) possess both glycosyltransferase and transpeptidase activities, and they work together with the SEDS-bPBP complexes to build the cell wall.^5^ Some peptidoglycan synthases are primarily devoted to synthesizing the cell wall along the cell periphery while others are dedicated to constructing the cell wall partition between daughter cells during cell division.^6^ Many studies have focused on characterizing these enzymes because of their importance in bacterial physiology and attractiveness as targets for antibiotic development. Indeed, the ß-lactam antibiotics that inhibit PBP transpeptidase activity have been one of the most clinically successful antibiotics to date.

**Figure 1:**
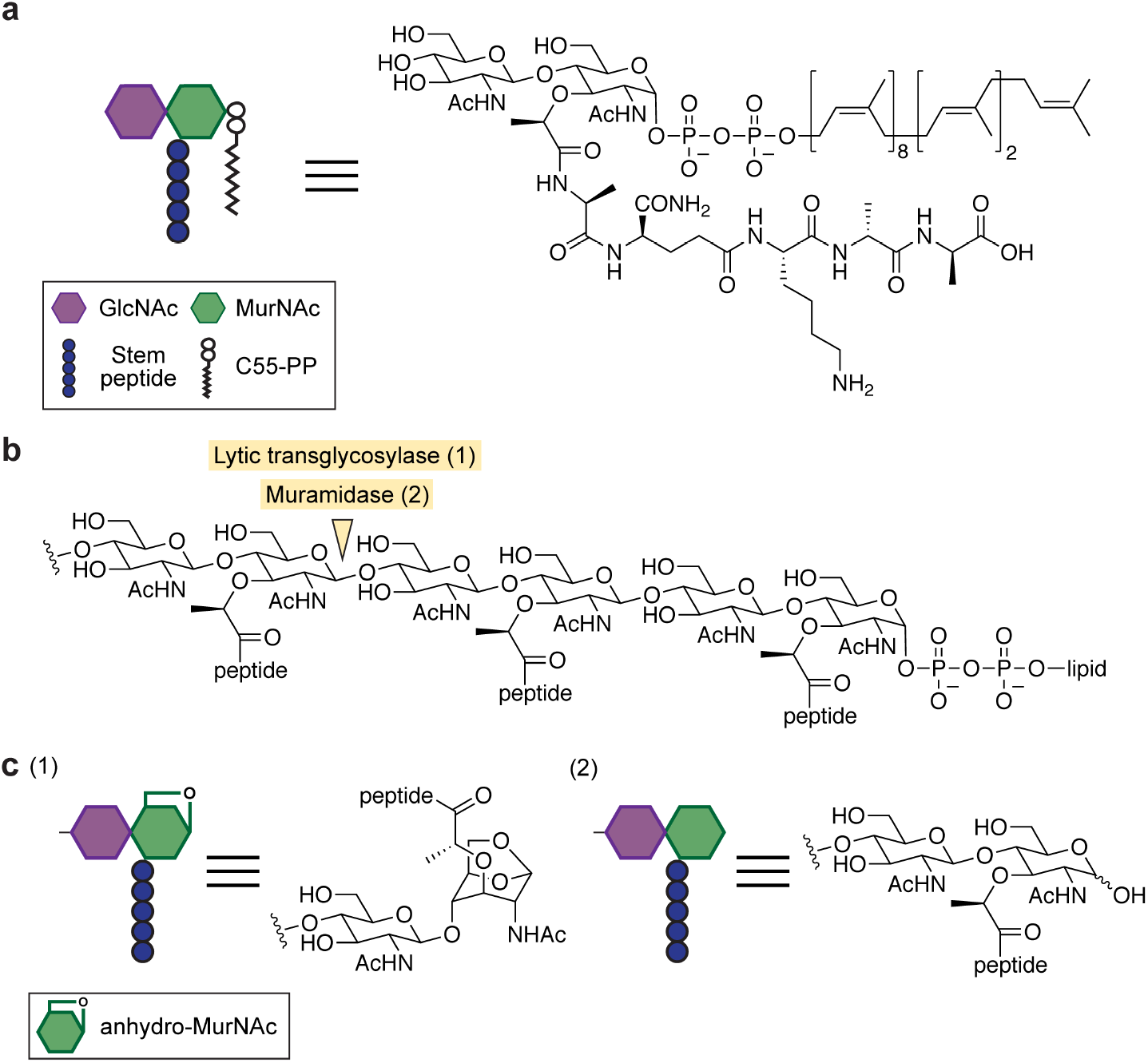
Lytic transglycosylases and muramidases cleave MurNAc(ß-1,4)GlcNAc bonds in peptidoglycan. **(a)** Structure of the peptidoglycan precursor Lipid II used in this study to make nascent peptidoglycan. **(b)** Structure of a peptidoglycan oligomer showing the chemical bond cleaved by lytic transglycosylases and muramidases. **(c)** Lytic transglycosylase and muramidase cleavage products have anhydro-MurNAc (1) and MurNAc (2) at the liberated reducing end, respectively.

Compared to peptidoglycan synthases, proteins responsible for peptidoglycan maturation have received relatively little attention. Integration of the newly synthesized peptidoglycan into the cell wall requires liberating the glycan strand from the lipid anchor that tethers it to the cytoplasmic membrane. One class of enzymes that have been implicated in this process are membrane-bound glycosidases, which are capable of cleaving the peptidoglycan sugar backbone. MltG is a cytoplasmic membrane-anchored lytic transglycosylase that has been proposed to fulfill this role by catalyzing a non-hydrolytic cleavage of the ß-1,4 glycosidic bond between N-acetylmuramic acid (MurNAc) and N-acetylglucosamine (GlcNAc) to yield muropeptide products containing an 1,6-anhydro-MurNAc (anhMurNAc) end (Figure 1b, 1c).^7,8^ MltG is a member of the YceG family proteins that are widely present in rod- and oval-shaped bacteria.^7^ Some bacteria, including *Staphylococcus aureus*, do not encode YceG family proteins, and other types of cell wall cleaving enzymes must liberate the glycan strand from the lipid anchor. In *S. aureus*, the membrane anchored N-acetylglucosaminidase SagB has been proposed to perform this role by forming a complex with a partner protein that controls cleavage site selection on the nascent peptidoglycan strand.^9^

Here, we have investigated two putative cell wall cleaving enzymes in *Streptococcus pneumoniae* that were previously reported to play roles in cell wall integrity. *S. pneumoniae* is an opportunistic pathogen that colonizes mucosa in the upper respiratory tract and can lead to a wide range of sometime fatal infections, including pneumonia, meningitis, and sepsis.^10^ As *S. pneumoniae* is increasingly resistant to clinically-used antibiotics, understanding its vulnerabilities is a priority.^11^ One protein studied here, SPD_1346 (formerly MltG; here renamed MpgA for <u>m</u>embrane-bound <u>p</u>eptidoglycan <u>g</u>lycosidase <u>A</u>) contains an extracellular YceG domain that resembles the lytic transglycosylase MltG.^7,12^ MpgA is essential in unencapsulated *S. pneumoniae* strains, and *mpgA*-null mutants can only be constructed in strains that also lack the bifunctional peptidoglycan synthase PBP1a.^13^ MpgA depletion causes spherical morphology, implying a role in cell shape.^13^ Based on these and other observations, it was proposed that MpgA’s putative lytic transglycosylase activity is crucial for peptidoglycan assembly along the cell periphery.^13,14^ The other protein studied here, SPD_0912 (formerly PMP23; renamed MpgB) shares structural homology with the *E. coli* lytic transglycosylase MltE and with a *B. subtilis* muramidase that hydrolyzes the ß-1,4 glycosidic bond between MurNAc and GlcNAc (Figure 1b, 1c).^15,16^ Although not essential, MpgB deletion causes mislocalization of division proteins and septal defects, suggesting that MpgB’s putative enzymatic activity is important in cell wall assembly at the septum.^17,18^ Here, we characterized the enzymatic functions of MpgA and MpgB, and found surprising differences from what we expected. We also identified a domain in MpgA that regulates where cleavage occurs along the glycan backbone. Our studies not only provide new insights into *S. pneumoniae*, but have broad implications for cell wall hydrolase function in other bacteria.

## RESULTS

*S. pneumoniae*, like other bacteria, contains multiple genes that encode confirmed or putative cell wall hydrolases. We focused on MpgA because it is essential and previous efforts to reconstitute its activity were unsuccessful.^13^ Most other cell wall hydrolases in *S. pneumoniae* are non-essential so we needed a strategy to differentiate those involved in cell wall assembly during normal growth from those that play other roles. In *E*.*coli* and *S. aureus*, ß-lactam hyper-sensitivity in deletion mutants has been used to identify cleavage enzymes that act early in the cell wall assembly pathway.^9,19–21^ Hyper-sensitivity is notable because deleting cell wall hydrolases often makes cells more rather than less tolerant of ß-lactam treatment, evidently because reducing cell wall degrading activity is advantageous when cell wall synthesis activity is inhibited.^22–25^ We individually deleted six known or putative cell wall hydrolase genes in *S. pneumoniae* and screened the mutants for hyper-susceptibility to oxacillin and methicillin. These ß-lactams preferentially target PBP2x, which is an essential bPBP required for cell division.^26^ In a spot titer assay, only Δ*mpgB* cells displayed increased ß-lactam susceptibility (Figure 2a, Supplementary Figure 1). Moreover, when grown in liquid culture, Δ*mpgB* cells lysed more rapidly than wildtype cells upon ß-lactam treatment (Figure 2b). The hyper-sensitivity phenotype was reversed by complementing Δ*mpgB* with ectopically expressed *S. pneumoniae mpgB* or with a *mpgB* homolog from *Streptococcus thermophilus, Bacillus subtilis* or *Enterococcus faecalis* (Figure 2c). These results suggested that the MpgB’s activity is important during nascent cell wall assembly in *S. pneumoniae*. Because literature evidence also supports an important role for MpgB in cell wall assembly, we examined it as well as MpgA.^18^

**Figure 2.**
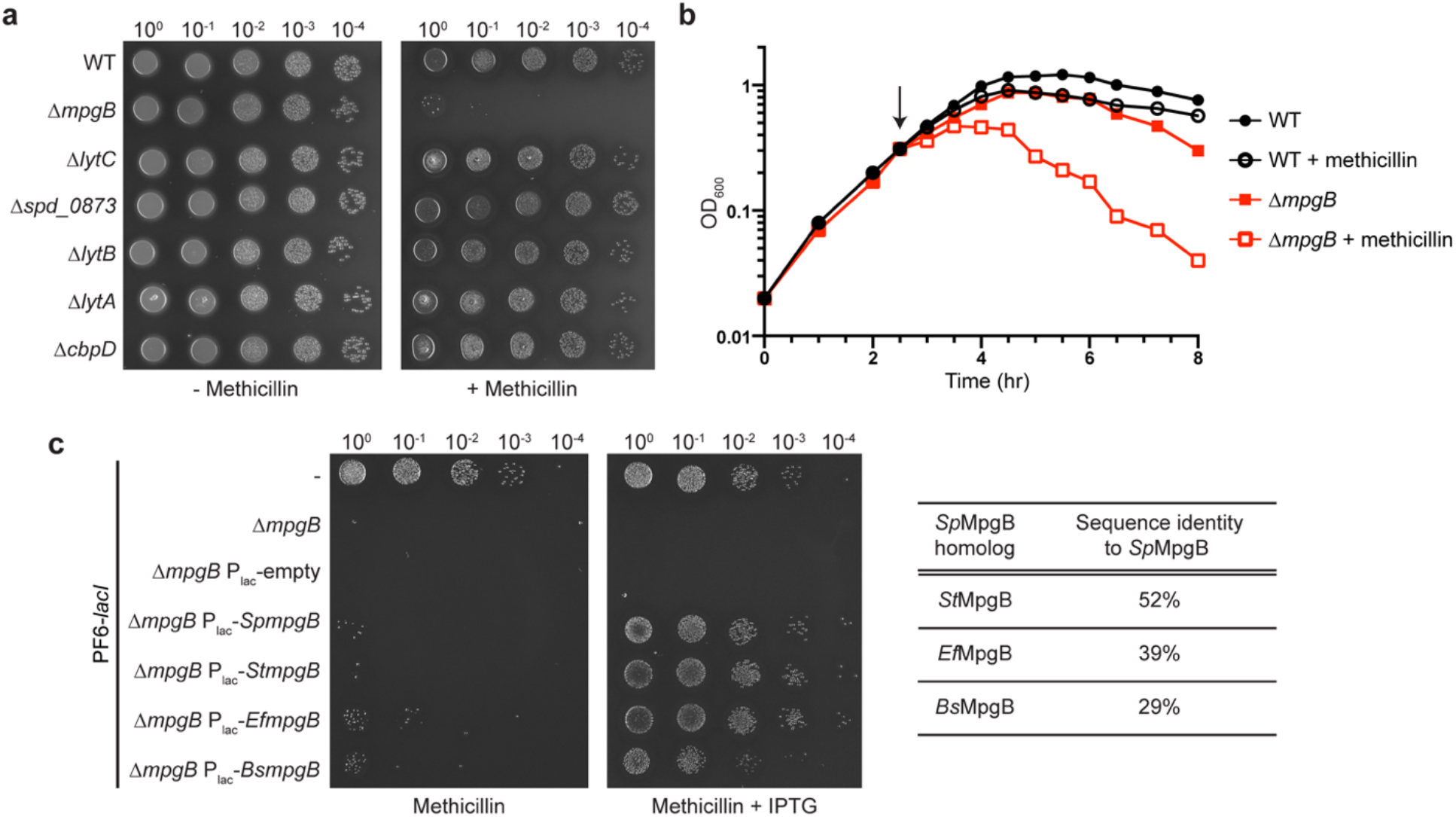
MpgB deletion renders *S. pneumoniae* hyper-sensitive to methicillin. **(a)** *S. pneumoniae* cells containing the indicated gene deletion were spotted on agar plates with methicillin (64 ng/mL) to identify hyper-sensitivity phenotypes. **(b)** Representative growth curves of *S. pneumoniae* wild-type and Δ*mpgB* strains showing effects of adding methicillin (black arrow; 16 ng/mL). **(c)** Methicillin sensitivity of Δ*mpgB* cells can be rescued by expressing MpgB homologs from *S. pneumoniae, S. thermophilus, E. faecalis* or *B. subtilis* under an IPTG-inducible promoter (P_lac_). Cells were spotted on agar containing methicillin (64 ng/mL) and IPTG (1 mM). Strains used in this experiment express LacI that prevents leaky expression of P_lac_.^41^

MpgA and MpgB were assumed to have enzymatic activity, but no direct evidence for enzymatic function has been reported. We therefore purified MpgA, MpgB, and several homologs from other organisms to test for activity (Supplementary Fig. 2). Only one of these proteins, the MpgA homolog *E. coli* MltG, which has lytic transglycosylase activity, has been biochemically characterized.^7^ We prepared substrates for our assays by polymerizing fluorescently-labeled Lipid II, the peptidoglycan precursor, using a polymerase engineered to make short peptidoglycan strands.^27^ The labeled oligomers were incubated with the purified proteins and the reaction products were separated by SDS-PAGE and subjected to in-gel visualization (Figure 3a). The untreated peptidoglycan oligomers run as a ladder where each band differs by a monomer (*i*.*e*., disaccharide) unit. In lanes treated with either MpgB or MpgA we observed clear changes in the banding pattern compared with the control (Figures 3b, 3c). For both proteins, mutating the proposed catalytic residue abolished reaction and we concluded that both proteins had glycosidase activity. However, there were striking differences in the products detected in the PAGE assay. Oligomers treated with MpgB were degraded to a predominant, rapidly-migrating band with a mobility corresponding to Lipid II, the lipid-linked disaccharide-peptide starting material. In contrast, oligomers treated with MpgA were degraded to a predominant band with a mobility corresponding to a product having seven disaccharide units (*i*.*e*., Lipid 14). These distinct cleavage products were also observed for homologs of MpgB and MpgA. Therefore, these two classes of enzymes have an intrinsic ability to cleave nascent peptidoglycan at different sites relative to the lipid anchor on the reducing end of the polymer.

**Figure 3.**
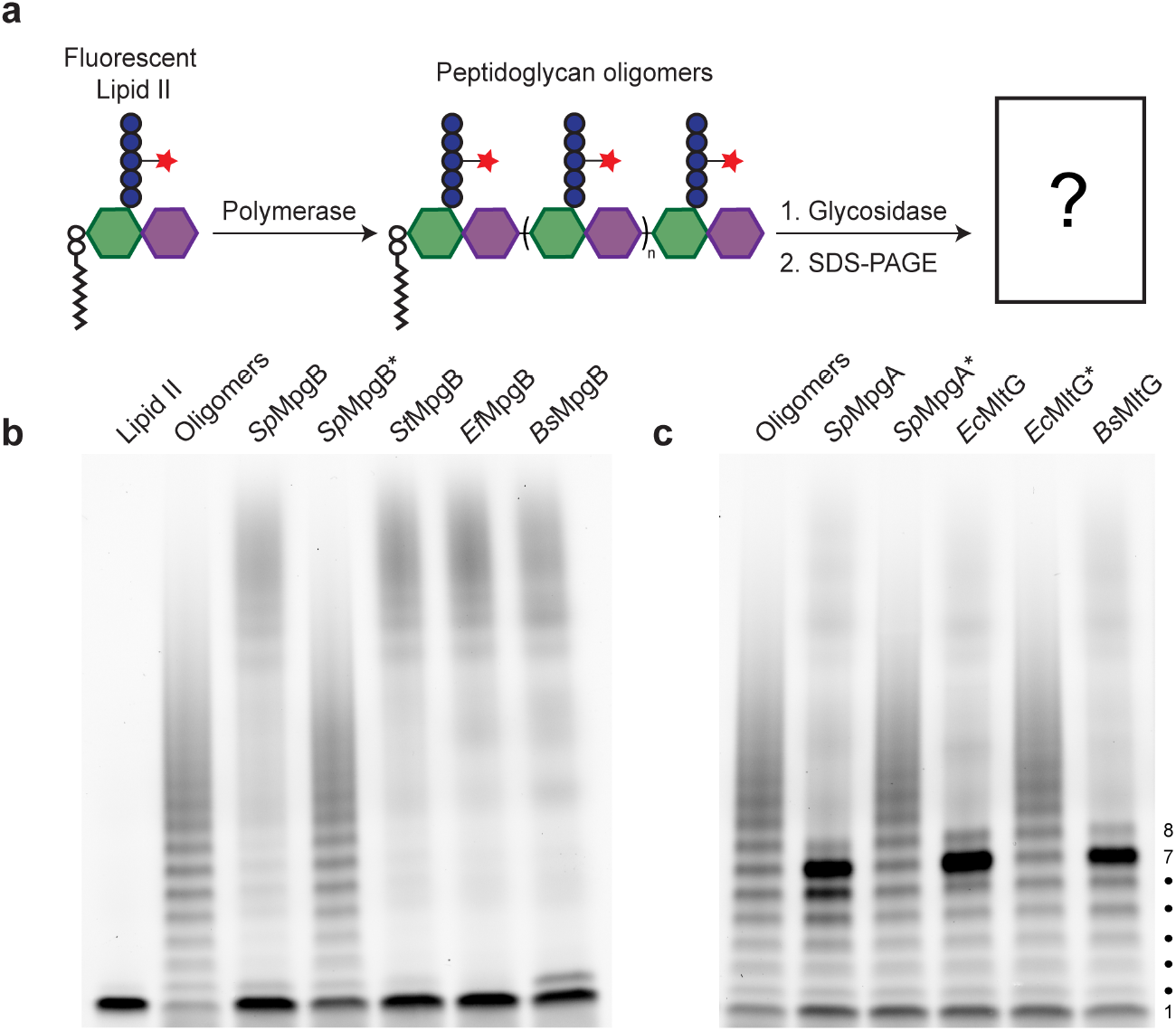
MpgA and MpgB cleave nascent peptidoglycan at different positions along the backbone. **(a)** Schematic of the assay used to detect peptidoglycan glycosidase activity. The fluorescent dye ATTO488 is attached to the stem peptide lysine for in-gel visualization of peptidoglycan products. **(b)** MpgB cleaves nascent peptidoglycan to Lipid II *in vitro* (bottom band, #1). **(c)** MpgA and other YceG family proteins cleave nascent peptidoglycan to a major lipid-linked product with a mobility corresponding to seven disaccharide repeats. In **b** and **c**, asterisks indicate catalytically inactive variants. Diffuse signal high in the gel in reaction lanes are products released from the lipid anchor by cleavage (see Figure 4).

Before addressing mechanisms for product length control, we wanted to establish the nature of the bond cleavage reactions. We observed a diffuse signal in the upper half of the lanes for the enzyme-treated samples. This signal is due to cleavage products that lack the diphospholipid anchor.^9^ We used liquid chromatography-mass spectrometry (LC-MS) to characterize these products. Unlabeled peptidoglycan oligomers were prepared, cleaved with MpgB or MpgA, and the reaction samples were treated with sodium borohydride (NaBH_4_) to reduce any hydrolysis products (Figure 4a: products A1-A4, Supplementary Fig 3 & 4). In the *Sp*MpgB sample, the mass of the major peak corresponded to a tetrasaccharide hydrolysis product, but we also detected small peaks for larger products (Figure 4b, panel 1). Addition of mutanolysin, a well-characterized muramidase, to the *Sp*MpgB-treated sample cleaved the products to the disaccharide-monopeptide monomer unit (Figure 4b, panel 2). *Sp*MpgB was previously speculated to have either muramidase activity or lytic transglycosylase activity, and our product analysis showed that it is a muramidase.^13,18^ *Bs*MpgB and *Ef*MgpB were also shown to have muramidase activity (Figure 4b, panels 3 and 4).

**Figure 4.**
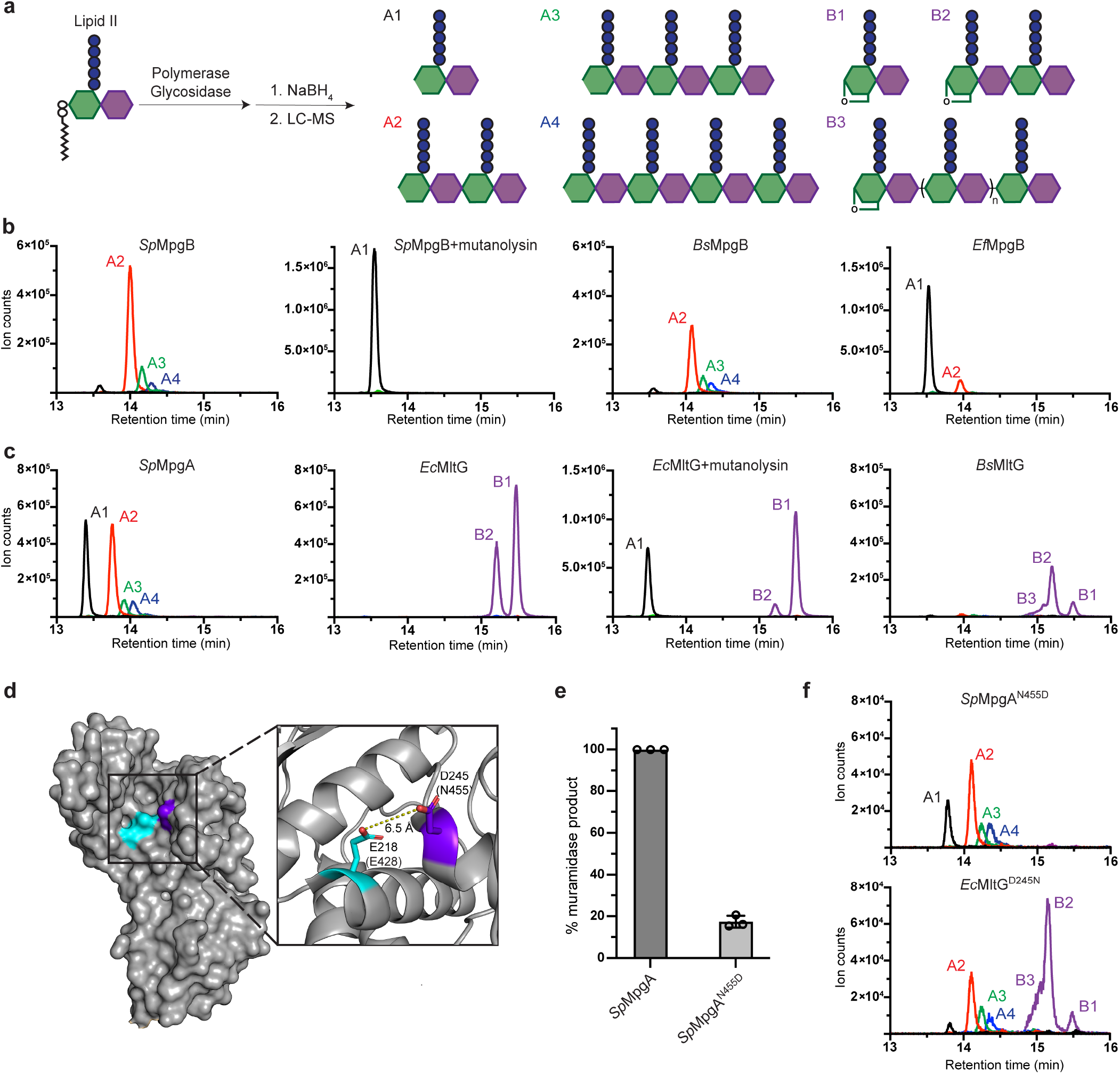
MpgA is a muramidase rather than a lytic transglycosylase like MltG. **(a)** Schematic of the LC-MS assay used to assess peptidoglycan cleavage activity. Products released from the lipid anchor by cleavage activity are detected in this assay. **(b)** MpgB is a peptidoglycan muramidase. **(c)** MpgA is a muramidase unlike its lytic transglycosylase homologs in *E. coli* and *B. subtilis*. For **b** and **c**, extracted ion chromatograms were generated by plotting the intensity of the signal corresponding to the mass-to-charge ratio (m/z) of the muramidase products (A1-A4) and lytic transglycosylase products (B1-B3). Lytic transglycosylase product lengths are distinguished by their charge states. B3 indicates unresolved lytic transglycosylase oligomer products having n>0. **d)** Structure of the *Ec*MltG active site with the catalytic residue, Glu218, and a highly conserved aspartate, Asp245, shown in cyan and purple, respectively (PDB: 2R1F). The corresponding residues in *Sp*MpgA are indicated in parentheses. **(e)** Normalized levels of muramidase products detected by LC-MS for *Sp*MpgA and the *Sp*MpgA^N455D^ variant. **(e)** LC-MS analysis of *Sp*MpgA^N455D^ and *Ec*MltG^D245N^ cleavage products shows muramidase activity for the latter. See Figure 4a for peak assignments.

The results for MpgA were more surprising because bioinformatics predicted it was a lytic transglycosylase.^7,13^ Lytic transglycosylases release muropeptides with anhMurNAc ends that cannot be reduced by NaBH_4_ (Figure 4a: products B1-B3, Supplementary Fig 3 & 4). However, the products for MpgA had the same masses as those for MpgB, showing that this protein also has muramidase activity (Figure 4c, panel 1). In contrast, and as expected, *Ec*MltG generated anhMurNAc products (Figure 4c, panel 2 & 3).^7^ We also confirmed that the *B. subtilis* homolog, *Bs*MltG, is a lytic transglycosylase (Figure 4c, panel 4).

Our discovery that MpgA is a muramidase rather than a lytic transglycosylase is consistent with previous studies reporting that anhydromuropeptide species are undetectable in the pneumococcal cell wall.^28^ However, the known lytic transglycosylase *Ec*MltG was able to complement a Δ*mpgA* deletion. Therefore, MpgA and *Ec*MltG were speculated to have the same enzymatic activity in cells, and failure to detect anhMurNAc in the *S. pneumoniae* cell wall was attributed to low abundance of these modifications.^13^ To verify that it would be possible to detect anhMurNAc species due to the action of MltG, we constructed a *S. pneumoniae* Δ*mpgA* strain that ectopically expressed either *Ec*MltG or *Sp*MpgA. We analyzed the muropeptides released from the isolated cell wall from these strains after mutanolysin digestion (Supplementary Fig. 5). In line with our biochemical results, muropeptides containing anhMurNAc ends were observed in peptidoglycan isolated from *Ec*MltG-expressing cells, but not from peptidoglycan isolated from *Sp*MpgA-expressing cells. These results support our conclusion that *Sp*MpgA, unlike its homolog *Ec*MltG, is a peptidoglycan muramidase.

The predicted structure of *Sp*MpgA is nearly identical to the determined structures of the YceG domains of *Ec*MltG and *Listeria monocytogenes* MltG, but our findings show that its bond-cleaving activity is different.^13^ To identify residues that might play a role in determining whether a YceG homolog functions as a lytic transglycosylase or a muramidase, we aligned sequences of >10,000 YceG domains to identify conserved residues (Supplementary Figure 6). Several highly conserved residues are replaced in *Sp*MpgA compared with verified YceG lytic transglycosylases, but *Sp*MpgA residue Asn455 particularly caught our attention because of its predicted proximity to the putative catalytic glutamate (Figure 4d).^7,12^ This residue is an aspartate in 95% of YceG-family proteins, including *Ec*MltG (Supplementary Figure 6). We wondered if the presence of Asn or Asp at this position was sufficient to determine if a YceG-family protein had muramidase or lytic transglycosylase activity. A N455D substitution of *Sp*MpgA reduced enzymatic activity, but did not result in anhMurNAc products (Figure 4e, 4f). However, replacing the corresponding aspartate with asparagine in *Ec*MltG (EcMltG^D245N^) led to the appearance of muramidase products (Figure 4f). Although other amino acids are involved in determining whether a MltG homolog is a lytic transglycosylase or a muramidase, we can conclude from the results on *Ec*MltG^D245N^ that the conserved active site Asn we identified in *Sp*MpgA indeed plays a role in determining muramidase activity. We therefore suggest that other YceG family proteins with an asparagine at this position, which are mainly found in *Streptococcus* and *Lactococcus* species, will have muramidase activity (Supplementary Figure 6). In showing that closely related cell wall cleaving enzymes can make different products, our findings highlight the need to corroborate any predicted enzymatic functions. We note that recently developed methods to make peptidoglycan *in vitro* from readily obtained Lipid II now make this straightforward.^29–31^

We next turned our attention to understanding length control of the lipid-linked cleavage products. Similar to mutanolysin, MpgB homologs cleave nascent peptidoglycan to Lipid II, but all the YceG proteins showed a predominant cleavage product with seven disaccharide repeats based on the gel mobility analysis. A notable difference between MpgB and the YceG proteins is that the latter contain a membrane-proximal LysM-like subdomain between the transmembrane helix and the catalytic domain (Figure 5a). LysM domains are widely distributed among peptidoglycan hydrolases and are proposed to modulate binding to peptidoglycan.^32,33^ Although they are typically found in hydrolases that bind to mature (crosslinked) cell wall, it would be reasonable to speculate that the membrane-proximal LysM-like domain in YceG proteins binds nascent peptidoglycan near its membrane anchor to help direct the peptidoglycan strand into the active site for cleavage. Based on the crystal structure, the distance from the LysM-like subdomain to the catalytic residue is approximately 50 °, but there are some additional residues between the LysM-like domain and the TM helix anchoring the protein in the membrane, which may extend the distance from the membrane to the active site.^34^ Each disaccharide unit in peptidoglycan spans approximately 10 °. Overall, the dimensions of MpgA are sufficiently well-matched with what we now know about its cleavage site preferences for the proposed model to be plausible. To test whether the LysM-like domain in *Sp*MpgA does indeed affect product length, we expressed and purified a *Sp*MpgA variant containing an internal deletion of this domain (*Sp*MpgA^ΔLysM^) and tested its activity. *Sp*MpgA^ΔLysM^ retained its muramidase activity, but released a lipid-linked cleavage product that was approximately half as long as the product formed when the LysM-like domain was present (mobility consistent with Lipid 8; Figure 5b, 5c). We conclude that the LysM-like domain is crucial for MpgA cleavage site selection, supporting the hypothesis that it binds nascent peptidoglycan near its membrane anchor.

**Figure 5.**
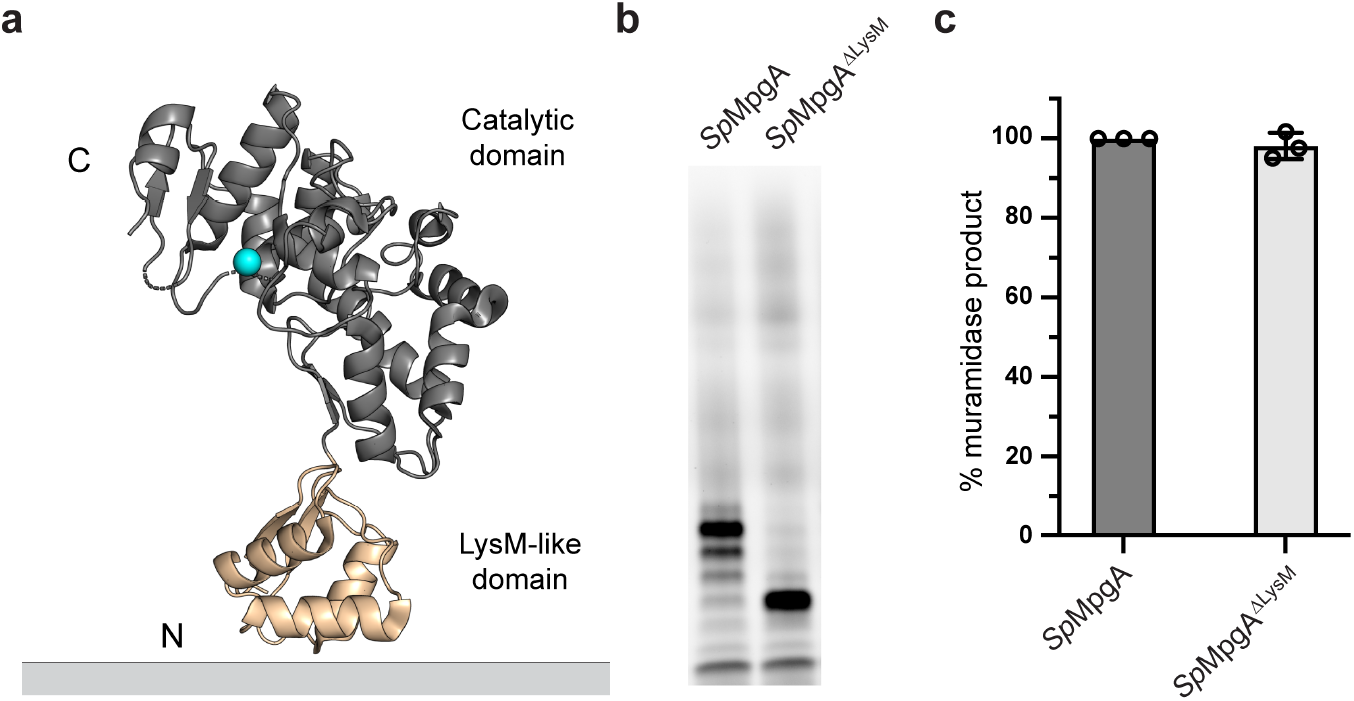
The LysM-like domain of MpgA controls the length of lipid-linked cleavage product. **(a)** The crystal structure of the *Listeria monocytogenes* MltG homolog (PDB: 4IIW) shows the N-terminal LysM-like subdomain (tan) and the catalytic subdomain (gray) with the catalytic glutamate in cyan. The LysM-like subdomain is connected to a transmembrane helix that anchors the protein to the cytoplasmic membrane. **(b)** Shorter lipid-linked cleavage products are generated when MpgA lacks the LysM-like domain (*Sp*MpgA^ΔLysM^). **(c)** Deletion of LysM-like domain does not affect the catalytic efficiency of MpgA.

## DISCUSSION

By reconstituting the activities of two *S. pneumoniae* membrane-bound peptidoglycan glycosidases and their homologs, we have made several new discoveries that are also relevant to related glycosidases in other organisms. First, we have shown that not all YceG family proteins are lytic transglycosylases: despite its resemblance to *Ec*MltG, MpgA is a muramidase. Second, the LysM-like subdomain in YceG family proteins confers on these proteins the ability to cleave the glycan backbone at a distance of ∼70 ° from the lipid anchor. We have also shown that MpgB, speculated by some to be a muramidase and by others to be a lytic transglycosylase, has muramidase activity. Unlike MpgA, MpgB does not contain another subdomain and it cleaves nascent peptidoglycan at the MurNAc-GlcNAc bond closest to the lipid anchor.

The studies reported here focus primarily on *in vitro* biochemistry of *S. pneumoniae* MpgA and MpgB, but it is worth asking what purpose these proteins serve in cells. Unlike MpgA and MpgB, many glycosidases do not localize in the cytoplasmic membrane; these other glycosidases tend to degrade mature peptidoglycan to enable daughter cell separation, facilitate cell wall recycling or remodel the cell wall.^35,36^ LysM domains are often found in these proteins and provide cell wall-targeting capability.^32^ Unlike proteins that degrade mature peptidoglycan, cell wall hydrolases anchored in the cytoplasmic membrane appear to act early in the peptidoglycan assembly pathway, cleaving uncrosslinked regions of newly-formed substrates that are not fully integrated into the cell wall matrix. For example, *S. aureus* LytH is a membrane-bound amidase that removes stem peptides only from uncrosslinked peptidoglycan.^21^ Stem peptide removal from nascent peptidoglycan controls cell size by limiting substrate availability and regulating spatial localization of peptidoglycan synthases. *E. coli* MltG is a membrane-bound lytic transglycosylase that was proposed to terminate elongation by cleaving nascent polymer as it emerges from the peptidoglycan synthase active site.^7^ Supporting a role in terminating elongation, the presence of MltG reduces glycan strand lengths in the *E. coli* cell wall.^7^ *S. aureus* SagB, a membrane-bound glucosaminidase, also reduces glycan strand length.^37^ Because *in vitro* studies showed that SagB cleaves nascent peptidoglycan *after* it dissociates from a peptidoglycan synthase, SagB was proposed to function as a “release factor” that cleaves partially crosslinked peptidoglycan polymers from their lipid anchor. This allows full integration of new strands into the mature cell wall while leaving a competent lipid-linked substrate in the membrane that can be extended once it rebinds to a peptidoglycan synthase.^9,38,39^

We favor the hypothesis that MpgA and MpgB, like SagB, serve as peptidoglycan release factors (Figure 6). These proteins are anchored to the membrane and cleave nascent peptidoglycan bonds that are found at a distance from the lipid anchor that is compatible with the estimated distance of their active sites from the membrane. We found that nascent peptidoglycan was an excellent substrate for both enzymes. Although we did not test whether MpgA and MpgB can also cleave crosslinked peptidoglycan, no cleavage products were detected in previous attempts to reconstitute these enzymes using isolated sacculi comprising crosslinked peptidoglycan as substrates.^13,18^

**Figure 6.**
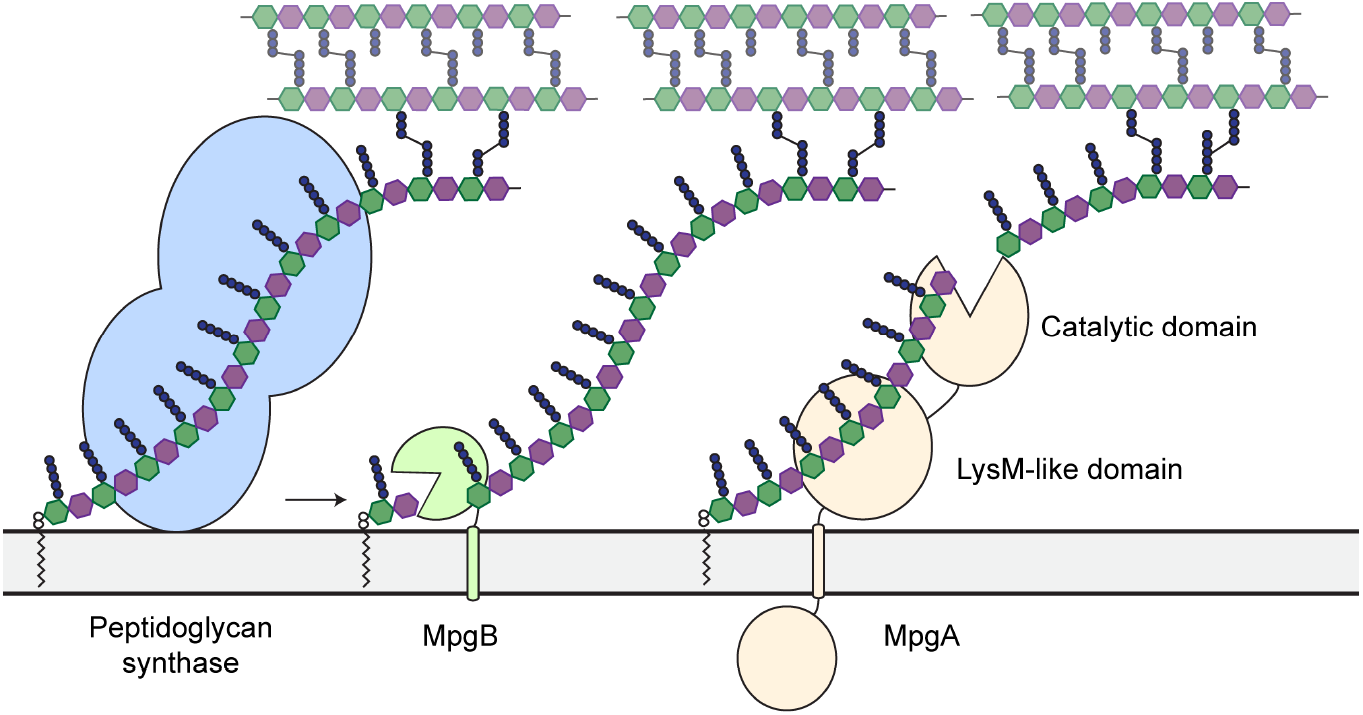
Working model for MpgA and MpgB function in cells. Peptidoglycan synthases polymerize Lipid II and crosslink the nascent glycan strand into the existing cell wall. MpgA and MpgB cleave nascent peptidoglycan at different distances from the lipid anchor to permit full integration of the released polymer into the cell wall. The lipid-linked products are substrates for further extension by peptidoglycan synthases.^9,38,39^

*S. aureus* SagB provides an interesting comparison to YceG family proteins. SagB forms a complex with SpdC, which is an integral membrane protein that shares some homology with the CAAX protease family that processes polyprenylated proteins.^9,40^ The SagB-SpdC complex has been shown to cleave nascent peptidoglycan at a defined distance from the lipid anchor, and it was proposed that SpdC binds the lipid anchor on nascent peptidoglycan and guides the glycan strand up into the SagB active site.^9^ SagB itself is a functional glucosaminidase without SpdC *in vitro*; however, like MpgB it cleaves nascent peptidoglycan to Lipid II rather than the lipid-linked oligomeric products observed when its partner protein is not present. SagB loses function in Δ*spdC* cells, suggesting that control over product length, and not simply enzymatic activity, is important.^9^ The role of the LysM-like domain in YceG family proteins mirrors SpdC in its ability to determine cleavage site selectivity by controlling the location of the glycosidase active site with respect to the membrane. It is unknown whether cleavage site control is important in YceG family proteins; however, an MltG variant lacking the LysM-like subdomain was not functional in *E. coli*.^34^ Because enzymatic activity for this MltG variant was not confirmed, it is not clear whether lack of function reflected an inability to cleave nascent peptidoglycan at all or was due to lack of wildtype cleavage site selectivity.

A fundamental question that is well beyond the scope of this study is how MpgA and MpgB coordinate their activities with peptidoglycan synthases and tailoring enzymes to build the *S. pneumoniae* cell wall. Previous genetic and cell biological studies indicate that MpgA and MpgB act primarily during cell elongation and division, respectively.^13,18^ Models that posit direct physical interactions of MpgA and MpgB with peptidoglycan synthases have been proposed and provide intuitively appealing mechanisms for how cleavage and synthase activities are coordinated; however, evidence to support the *functional* importance of such interactions is lacking.^7,18^ The observation that MpgA and MpgB can be complemented with homologs sharing limited sequence identity strongly suggests that the enzymatic activities of these glycosidases determine their effects in cells. The enzymatic activities are conserved, but protein-protein interfaces undergo co-evolution and often are not conserved between species. It is possible that activities of synthases and membrane-anchored hydrolases are “matched” through less elegant mechanisms, including relative expression levels of proteins, protein localization, and substrate availability. Peptidoglycan tailoring modifications installed by other enzymes may also modulate availability of cleavage sites. With a clearer picture emerging for the biochemistry of membrane-bound cell wall hydrolases, we can begin to consider how their activities are coordinated with activities of synthetic and tailoring enzymes to build new cell wall.

## Acknowledgements

We thank Tyler Sisley for assistance with the initial characterization of *Sp*MpgB and Julia Page for critical reading of the manuscript. We also thank Genevieve Dobihal and David Rudner for the kind gift of *B. subtilis* PY79 genomic DNA. LC-MS data was acquired on an Agilent 6520 Q-TOF mass spectrometer supported by the Taplin Funds for Discovery Program. Funding for this work was provided by National Institutes of Health grants R01 AI139011 and R01 AI148752 to S.W. A.T. is supported in part by the Funai Overseas Scholarship.

## Author Contributions

A.T. and S.W. conceived the project. Experiments were designed and performed by A.T. Data was analyzed by A.T. and S.W. The manuscript was written by A.T. and S.W.

## Competing Interests

The authors declare no competing interest.

## Methods

### Materials

Unless otherwise indicated, all chemicals and reagents were purchased from Sigma-Aldrich. Restriction enzymes were purchased from New England Biolabs. Oligonucleotide primers were purchased from Integrated DNA Technologies. Culture media were purchased from Becton Dickinson. *S. thermophilus* LMG 18311 genomic DNA (gDNA) was purchased from ATCC (ATCC BAA-250D-5). *S. pneumoniae* Δ*murMN* Lipid II was isolated from cells as described previously^29,30^. *S. aureus* SgtB and SgtB^Y181D^ were expressed and purified as previously reported^27,42^.

### Bacterial strains, plasmids, oligonucleotide primers and culture conditions

*E. coli* strains were grown with shaking at 37°C in lysogeny broth (LB), Terrific Broth (TB) or on agarized LB plates with appropriate additives. *S. pneumoniae* strains were cultured statically in Todd Hewitt broth containing 0.5% yeast extract (THY) at 37°C in an atmosphere containing 5% CO_2_. Pre-poured Trypticase Soy Agar with 5% Sheep Blood (TSAII 5%SB) plates with a 5 mL overlay of 1% nutrient broth agar or TSA plates containing 5% defibrinated sheep blood with appropriate additives were used for *S. pneumoniae* when growth on solid media was required. The following concentration of antibiotics were used: carbenicillin, 50 µg/mL; chloramphenicol, 35 µg/mL; erythromycin, 0.2 µg/mL; gentamycin, 100 µg/mL; kanamycin, 50 µg/mL (*E. coli*) or 250 µg/mL (*S. pneumoniae*); spectinomycin, 200 µg/mL. The bacterial strains, plasmids and oligonucleotide primers used in this study are summarized in Supplementary Tables. Protocol for plasmid construction can be found in the Supplementary Methods.

### Protein expression: general procedure

A previously described protocol was used for protein expression.^3^ *E. coli* C43(DE3) containing the expression plasmid(s) of interest was grown in 1 L TB supplemented with the appropriate antibiotics at 37°C with shaking until OD_600_ ∼0.7. The culture was cooled to 20°C and protein expression was induced by adding 500 µM isopropyl β-D-1-thiogalactopyranoside (IPTG) (His_6_-tagged protein) or 500 µM IPTG and 0.1% arabinose (FLAG-tagged protein). Cells were harvested 18 h post-induction by centrifugation (4,200 x g, 15 min, 4 °C) and the pellet was stored at −80 °C.

### Purification of His_6_-tagged proteins

All steps after cell lysis were performed at 4°C. Cells were resuspended in 50 mL lysis buffer (50 mM HEPES pH 7.5, 150 mM NaCl) and lysed by passaging the resuspended cells through a cell disruptor (EmulsiFlex-C5, Avestin) at 15,000 psi three times. Cell debris was removed by centrifugation (12,000 x g, 5 min, 4 °C) and the membrane fraction was collected by ultracentrifugation of the supernatant (100,000 x g, 1 h, 4 °C). The membrane pellet was resuspended in 30 mL solubilization buffer (50 mM HEPES pH 7.5, 500 mM NaCl, 1% n-dodecyl β-D-maltoside (DDM), 10% glycerol) using a glass dounce tissue grinder (Wheaton). The resulting mixture was stirred for 1 h at 4°C before ultracentrifugation (100,000 x g, 30 min, 4 °C). The resulting supernatant was supplemented with 0.75 mL pre-equilibrated Ni-NTA resin (Qiagen) and 20 mM imidazole and stirred for 30 min at 4 °C. The sample was then loaded onto a gravity column and washed with 30 mL buffer A (50 mM HEPES pH 7.5, 500 mM NaCl, 0.05% DDM, 10% glycerol) containing 20 mM imidazole and 30 mL buffer A containing 40 mM imidazole. The protein was then eluted in 10 mL buffer A containing 300 mM imidazole. The eluate was further purified by size exclusion chromatography (SEC) with a Superdex 200 10/300 GL column (MpgB) or a Superose 6 Increase 10/300 GL column (MpgA/MltG) equilibrated in buffer A. Fractions containing the target protein were concentrated by centrifugal filtration. The absorbance at 280 nm was measured using a NanoDrop One/One Microvolume UV-Vis Spectrophotometer (ThermoFisher Scientific) and the predicted extinction coefficient was used to calculate concentration. Protein samples were then aliquoted and stored at −80 °C.

### Purification of FLAG-tagged proteins

FLAG-tagged MpgBs were purified via the same protocol as His_6_-tagged proteins, above, with the following modifications. The supernatant containing the DDM-solubilized protein was supplemented with 2 mM CaCl_2_ and loaded onto a homemade 1 mL M1 α-Flag antibody resin. The resin was washed with 60 mL of buffer A supplemented with 2 mM CaCl_2_ and the bound protein was eluted from the column with 10 mL of buffer A supplemented with 5 mM EDTA and 0.2 mg/mL FLAG peptide (Genscript).

### Preparation of ATTO488-labeled Lipid II

*S. pneumoniae* Δ*murMN* Lipid II was fluorescently labeled with ATTO488 NHS-ester (ATTO-TEC) (NHS-ester:Lipid II = 8:1 mole ratio) in labeling buffer (0.1 M sodium borate pH 8.5) for 1 h at room temperature in the dark. The reaction mixture was then loaded onto a 1 mL Bakerbond C18 SPE column (J.T.Baker) by gravity flow. The column was sequentially washed with 1.5 mL 10%, 20%, 40% and 60% methanol to remove excess NHS-ester, and the labeled product was eluted with 100% methanol. The elution fraction was dried *in vacuo* and stored at −20°C until further use.

### In-gel detection of peptidoglycan glycosidase activity

Fluorescent peptidoglycan oligomers were prepared by incubating 0.25 µM ATTO488-labeled Lipid II with 1 µM SgtB^Y181D^ (monofunctional peptidoglycan glycosyltransferase with impaired processivity)^27^ in a 1x reaction buffer (50 mM Tris pH 7.0, 10 mM CaCl_2_ and 10% DMSO) at 30°C for 1 h. The polymerization reaction was heat-quenched at 95°C for 5 min. After cooling, the digestion reaction was set up by adding 1 µL of 50 µM glycosidase to 9 µL of the polymerization reaction product (total volume 10 µL). After incubating this mixture at 30°C for 3 h (MpgA/MltG) or 10 h (MpgB), the reaction was quenched by adding 10 µL 2x Laemmli sample buffer (Bio-Rad). The samples were then loaded onto a 4-20% Mini-PROTEAN TGX Precast Protein gel (Bio-Rad) and run at 180V. The gels were imaged using a Typhoon FLA 7000 imager.

### LC-MS analysis of peptidoglycan glycosidase products

A previously published method was adapted to characterize peptidoglycan glycosidase products.^3,30^ *S. pneumoniae* Lipid II (50 µM) and glycosidase (5 µM) were incubated with SgtB (2 µM: for reactions containing MpgA/MltG) or SgtB^Y181D^ (5 µM: for reactions containing MpgB) in 20 µL 1x reaction buffer (20 mM MES pH 6.5, 10 mM CaCl_2_ and 10% DMSO) at 30°C for 16 h. This reaction was heat-quenched at 95°C for 5 min and cooled to room temperature. Several glycosidase-treated samples were further digested by adding 3 µL mutanolysin (from *Streptomyces globisporus*: 4000 U/mL) and incubating at 37°C for 3 h. To reduce the muropeptide products, 50 µL NaBH_4_ (10 mg/mL) was added and the mixture was incubated at room temperature for 30 min. The pH was adjusted to ∼4 with 20% phosphoric acid and the samples were lyophilized to dryness overnight. Dried samples were resuspended in 18 µL H_2_O and analyzed by LC-MS on an Agilent Technologies 1200 series HPLC system in line with an Agilent 6520 Q-TOF mass spectrometer with ESI-MS operating in positive mode. Products were separated on a Waters Symmetry Shield RP18 column (5 µm; 3.9 mm x 150 mm) with a matching column guard using the following method: flow rate = 0.5 mL/min, 100% solvent A (H_2_O, 0.1% formic acid) for 5 min followed by a linear gradient of solvent B (acetonitrile, 0.1% formic acid) from 0-40% over 25 min. Agilent MassHunter Workstation Qualitative Analysis software was used for analyzing the MS data.

### *S. pneumoniae* strain construction

*S. pneumoniae* D39 Δ*cps* and its derivatives were transformed using a previously reported protocol.^43^ All deletion strains except Δ*mpgB* were constructed by transforming an antibiotic resistance cassette flanked by ∼1kb upstream and downstream regions of the target gene. A markerless Δ*mpgB* strain was constructed by first transforming a DNA cassette that includes an antibiotic resistance marker and a *B. subtilis sacB* gene flanked by ∼1kb upstream and downstream regions of *mpgB*. After antibiotic selection, a PCR fragment containing the upstream and downstream regions of *mpgB* was transformed and transformants were selected with 10% sucrose.^44^ Detailed protocols for strain construction can be found in the Supplementary Methods.

### Spot dilution assay and growth assessment of *S. pneumoniae* strains

Cultures were prepared by inoculating several colonies from each strain in 5 mL THY. For spot dilution assays, cells in mid-log phase were collected by centrifugation and normalized to OD_600_ = 1.0. The normalized cultures were serially diluted and 3 µL of each dilution was spotted onto TSA 5% SB plates containing the indicated additives. For growth assessment, cultures in mid-log phase were diluted to OD = 0.02 and growth was monitored by measuring OD_600_ at the indicated time points. Growth curves presented in the figure are representatives of three independent experiments.

### Digestion and LC-MS analysis of the *S. pneumoniae* cell wall

A previously reported protocol was adapted for isolating *S. pneumoniae* peptidoglycan.^45^ *S. pneumoniae* cultures were prepared in 20 mL THY and grown until OD_600_ ∼0.7. Cells were pelleted and resuspended in 1 mL 100 mM Tris pH 6.8/0.25% sodium dodecyl sulfate. After boiling the cells at 100°C for 30 min, the suspension was pelleted and washed 4x with 1 mL H_2_O. The washed pellet was resuspended in 1 mL 100 mM Tris Ph 6.8 containing 10 mM CaCl_2_, 10 mM MgCl_2_and 50 µg/mL trypsin and incubated at 37°C for 1 h before heat-inactivating the enzymes at 95°C for 5 min. The suspension was centrifuged at 21,000 x g for 3 min and the resulting pellet was washed once with 1 mL H_2_O. The pellet was then suspended in 500 µL 1 M HCl and incubated at 37°C for 4 h. The suspension was pelleted and washed 4x with 1 mL H_2_O. The resulting peptidoglycan pellet was suspended in 100 µL 12.5 mM NaHPO_4_ pH 5.5 and 10 µL mutanolysin (4000 U/mL) was added to this suspension. The digestion reaction proceeded at 37°C for 16 h. After heat-inactivating mutanolysin at 95°C for 5 min, the insoluble material was pelleted by centrifugation at 21,000 x g for 3 min. The supernatant was transferred to a new microcentrifuge tube and muropeptide products were reduced by adding 50 µL NaBH_4_ (20 mg/mL) and incubating at room temperature for 30 min. The pH was adjusted to ∼4 with 20% phosphoric acid before LC-MS analysis.

LC-MS analysis of the cell wall muropeptide products was conducted with the same instrument setup as described above using the following method: flow rate = 0.5 mL/min, 100% solvent A (H_2_O, 0.1% formic acid) for 5 min followed by a linear gradient of solvent B (acetonitrile, 0.1% formic acid) from 0-20% over 120 min.

## Notes

### Competing Interest Statement

The authors have declared no competing interest.

## References

1. Silhavy, T. J., Kahne, D. & Walker, S. The bacterial cell envelope. Cold Spring Harbor Perspectives in Biology 2, 1–17 (2010).

2. Meeske, A. J. et al. SEDS proteins are a widespread family of bacterial cell wall polymerases. Nature 537, 634–638 (2016).

3. Taguchi, A. et al. FtsW is a peptidoglycan polymerase that is functional only in complex with its cognate penicillin-binding protein. Nat. Microbiol. 4, 587–594 (2019).

4. Sjodt, M. et al. Structural coordination of polymerization and crosslinking by a SEDS–bPBP peptidoglycan synthase complex. Nat. Microbiol. 5, 813–820 (2020).

5. Cho, H. et al. Bacterial cell wall biogenesis is mediated by SEDS and PBP polymerase families functioning semi-autonomously. Nat. Microbiol. 1, 16172 (2016).

6. Massidda, O., Nováková, L. & Vollmer, W. From models to pathogens: How much have we learned about Streptococcus pneumoniae cell division? Environ. Microbiol. 15, 3133–3157 (2013).

7. Yunck, R., Cho, H. & Bernhardt, T. G. Identification of MltG as a potential terminase for peptidoglycan polymerization in bacteria. Mol. Microbiol. 99, 700–718 (2016).

8. Dik, D. A., Marous, D. R., Fisher, J. F. & Mobashery, S. Lytic transglycosylases: concinnity in concision of the bacterial cell wall. Crit. Rev. Biochem. Mol. Biol. 52, 503–542 (2017).

9. Schaefer, K. et al. Structure and reconstitution of a hydrolase complex that may release peptidoglycan from the membrane after polymerization. Nat. Microbiol. 6, 34–43 (2021).

10. Henriques-Normark, B. & Tuomanen, E. I. The pneumococcus: Epidemiology, microbiology, and pathogenesis. Cold Spring Harb. Perspect. Med. 3, 1–15 (2013).

11. Kim, L., McGee, L., Tomczyk, S. & Beall, B. Biological and epidemiological features of antibiotic-resistant Streptococcus pneumoniae in pre- and post-conjugate vaccine eras: A United States perspective. Clin. Microbiol. Rev. 29, 525–552 (2016).

12. Lee, M. et al. From Genome to Proteome to Elucidation of Reactions for All Eleven Known Lytic Transglycosylases from Pseudomonas aeruginosa. Angew. Chemie - Int. Ed. 56, 2735–2739 (2017).

13. Tsui, H. C. T. et al. Suppression of a deletion mutation in the gene encoding essential PBP2b reveals a new lytic transglycosylase involved in peripheral peptidoglycan synthesis in Streptococcus pneumoniae D39. Mol. Microbiol. 100, 1039–1065 (2016).

14. Stamsås, G. A. et al. Identification of EloR (Spr1851) as a regulator of cell elongation in Streptococcus pneumoniae. Mol. Microbiol. 105, 954–967 (2017).

15. Xu, Q. et al. Structures of a bifunctional cell wall hydrolase CwlT containing a novel bacterial lysozyme and an NlpC/P60 dl-endopeptidase. J. Mol. Biol. 426, 169–184 (2014).

16. Fukushima, T. et al. Identification and characterization of novel cell wall hydrolase CwlT: A two-domain autolysin exhibiting N-acetylmuramidase and DL-endopeptidase activities. J. Biol. Chem. 283, 11117–11125 (2008).

17. Pagliero, E. et al. The Inactivation of a New Peptidoglycan Hydrolase Pmp23 Leads to Abnormal Septum Formation in Streptococcus pneumoniae. Open Microbiol. J. 2, 107–114 (2008).

18. Jacq, M. et al. The cell wall hydrolase Pmp23 is important for assembly and stability of the division ring in Streptococcus pneumoniae. Sci. Rep. 8, 7591 (2018).

19. Templin, M. F., Edwards, D. H. & Holtje, J. V. A murein hydrolase is the specific target of bulgecin in Escherichia coli. J. Biol. Chem. 267, 20039–20043 (1992).

20. Cho, H., Uehara, T. & Bernhardt, T. G. Beta-lactam antibiotics induce a lethal malfunctioning of the bacterial cell wall synthesis machinery. Cell 159, 1310–1311 (2014).

21. Do, T. et al. Staphylococcus aureus cell growth and division are regulated by an amidase that trims peptides from uncrosslinked peptidoglycan. Nat. Microbiol. 5, 291–303 (2020).

22. Tomasz, A., Albino, A. & Zanati, E. Multiple antibiotic resistance in a Bacterium with suppressed autolytic System. Nature 227, 138–140 (1970).

23. Heidrich, C., Ursinus, A., Berger, J., Schwarz, H. & Höltje, J. V. Effects of multiple deletions of murein hydrolases on viability, septum cleavage, and sensitivity to large toxic molecules in Escherichia coli. J. Bacteriol. 184, 6093–6099 (2002).

24. Chung, H. S., Yao, Z., Goehring, N. W., Kishony, R. & Kahne, J. B. Rapid β-lactam-induced lysis requires successful assembly of the cell division machinery. Proc. Natl. Acad. Sci. U. S. A. 106, 21872–21877 (2009).

25. Flores-Kim, J., Dobihal, G. S., Fenton, A., Rudner, D. Z. & Bernhardt, T. G. A switch in surface polymer biogenesis triggers growth-phase-dependent and antibiotic-induced bacteriolysis. Elife 8, 1–23 (2019).

26. Kocaoglu, O., Tsui, H. C. T., Winkler, M. E. & Carlson, E. E. Profiling of β-lactam selectivity for penicillin-binding proteins in Streptococcus pneumoniae D39. Antimicrob. Agents Chemother. 59, 3548–3555 (2015).

27. Rebets, Y. et al. Moenomycin resistance mutations in Staphylococcus aureus reduce peptidoglycan chain length and cause aberrant cell division. ACS Chem. Biol. 9, 459–467 (2014).

28. Bui, N. K. et al. Isolation and analysis of cell wall components from Streptococcus pneumoniae. Anal. Biochem. 421, 657–666 (2012).

29. Qiao, Y. et al. Lipid II overproduction allows direct assay of transpeptidase inhibition by β-lactams. Nat. Chem. Biol. 13, 793–798 (2017).

30. Welsh, M. A. et al. Identification of a Functionally Unique Family of Penicillin-Binding Proteins. J. Am. Chem. Soc. 139, 17727–17730 (2017).

31. Schaefer, K., Owens, T. W., Kahne, D. & Walker, S. Substrate Preferences Establish the Order of Cell Wall Assembly in Staphylococcus aureus. J. Am. Chem. Soc. 140, 2442–2445 (2018).

32. Buist, G., Steen, A., Kok, J. & Kuipers, O. P. LysM, a widely distributed protein motif for binding to (peptido)glycans. Mol. Microbiol. 68, 838–847 (2008).

33. Mesnage, S. et al. Molecular basis for bacterial peptidoglycan recognition by LysM domains. Nat. Commun. 5, 4269 (2014).

34. Bohrhunter, J. L., Rohs, P. D. A., Torres, G., Yunck, R. & Bernhardt, T. G. MltG activity antagonizes cell wall synthesis by both types of peptidoglycan polymerases in Escherichia coli. Mol. Microbiol. mmi.14660 (2020). doi:10.1111/mmi.14660

35. Do, T., Page, J. E. & Walker, S. Uncovering the activities, biological roles, and regulation of bacterial cell wall hydrolases and tailoring enzymes. J. Biol. Chem. 295, 3347–3361 (2020).

36. Vollmer, W., Joris, B., Charlier, P. & Foster, S. Bacterial peptidoglycan (murein) hydrolases. FEMS Microbiol. Rev. 32, 259–286 (2008).

37. Chan, Y. G. Y., Frankel, M. B., Missiakas, D. & Schneewind, O. SagB glucosaminidase is a determinant of Staphylococcus aureus glycan chain length, antibiotic susceptibility, and protein secretion. J. Bacteriol. 198, 1123–1136 (2016).

38. Perlstein, D. L., Zhang, Y., Wang, T. S., Kahne, D. E. & Walker, S. The direction of glycan chain elongation by peptidoglycan glycosyltransferases. J. Am. Chem. Soc. 129, 12674–12675 (2007).

39. Welsh, M. A., Schaefer, K., Taguchi, A., Kahne, D. & Walker, S. Direction of Chain Growth and Substrate Preferences of Shape, Elongation, Division, and Sporulation-Family Peptidoglycan Glycosyltransferases. J. Am. Chem. Soc. 141, 12994–12997 (2019).

40. Manolaridis, I. et al. Mechanism of farnesylated CAAX protein processing by the intramembrane protease Rce1. Nature 504, 301–305 (2013).

41. Keller, L. E., Rueff, A.-S., Kurushima, J. & Veening, J.-W. Three New Integration Vectors and Fluorescent Proteins for Use in the Opportunistic Human Pathogen Streptococcus pneumoniae. Genes (Basel). 10, 394 (2019).

42. Wang, T. S. A. et al. Primer preactivation of peptidoglycan polymerases. J. Am. Chem. Soc. 133, 8528–8530 (2011).

43. Fenton, A. K., Mortaji, L. El, Lau, D. T. C., Rudner, D. Z. & Bernhardt, T. G. CozE is a member of the MreCD complex that directs cell elongation in Streptococcus pneumoniae. Nat. Microbiol. 2, 16237 (2016).

44. Li, Y., Thompson, C. M. & Lipsitch, M. A modified Janus cassette (sweet Janus) to improve allelic replacement efficiency by high-stringency negative selection in Streptococcus pneumoniae. PLoS One 9, 2–7 (2014).

45. Kühner, D., Stahl, M., Demircioglu, D. D. & Bertsche, U. From cells to muropeptide structures in 578 24 h: Peptidoglycan mapping by UPLC-MS. Sci. Rep. 4, 1–7 (2014).

