## Supplementary Information for "Biochemical reconstitution defines new functions for membrane-bound glycosidases in assembly of the bacterial cell wall"

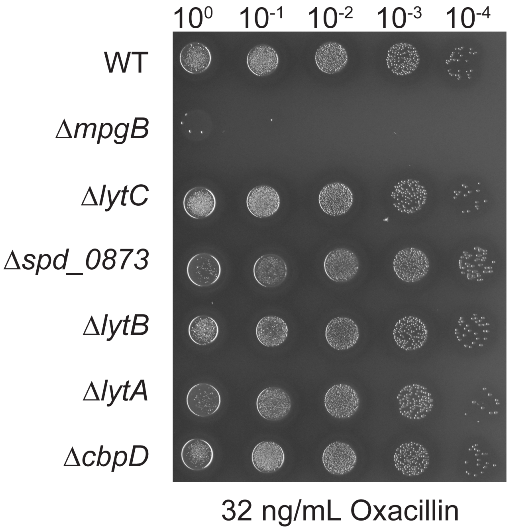


**Supplementary Figure 1. Δ*mpgB* cells are hypersensitive to oxacillin.** Spot dilution series of *S. pneumoniae* strains with the indicated deletion were plated on agar containing 32 ng/mL oxacillin.

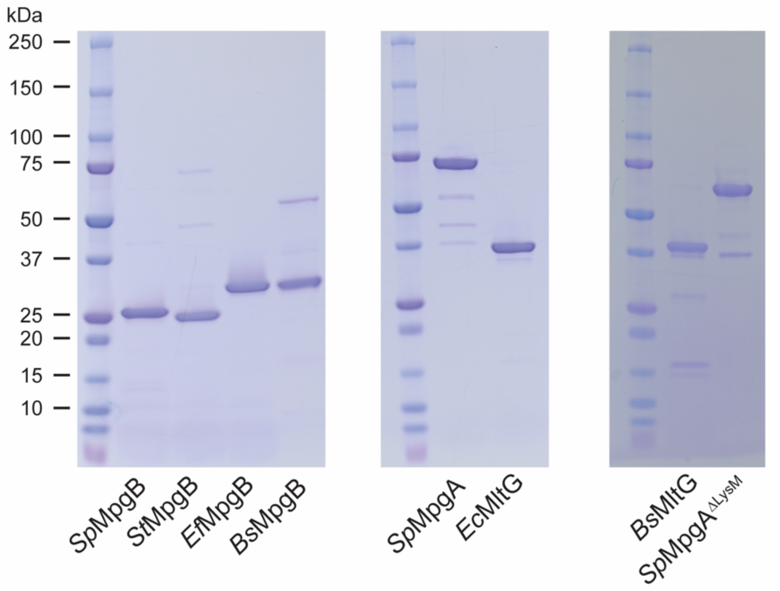


**Supplementary Figure 2. Coomassie stained gels of purified glycosidases used in this study.** ~2 μg protein was loaded per lane.

*Sp* = *S. pneumoniae*; *St* = *S. thermophilus*; *Ef* = *E. faecalis*; *Bs* = *B. subtilis*; *Ec* = *E. coli*

*Sp*MpgA$^{\Delta LysM}$ = *Sp*MpgA$^{\Delta D219\text{-}P295}$

A1

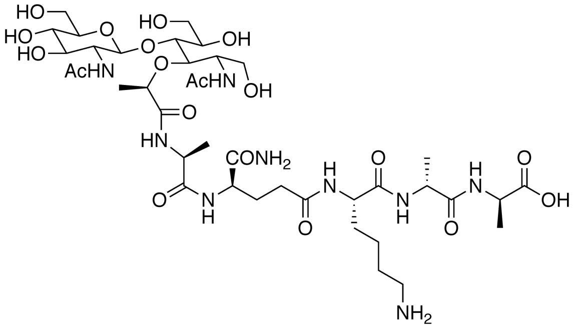

B1

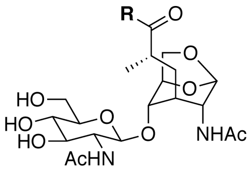

A2 (n = 0)
A3 (n = 1)
A4 (n = 2)

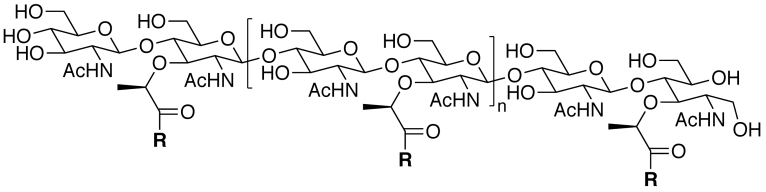

B2 (n = 0)
B3 (n ≥ 1)

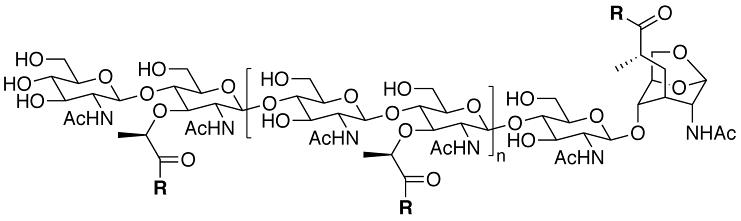

**R** = L-Ala-D-*i*Gln-L-Lys-D-Ala-D-Ala

**Supplementary Figure 3. Chemical structures of muropeptide products detected in digestion reactions following sodium borohydride reduction.** A1-A4 are peptidoglycan muramidase products and B1-B3 are lytic transglycosylase products with a 1,6-anhydroMurNAc end. See Figure 4a for the schematic of LC-MS assay used to detect cleavage products.

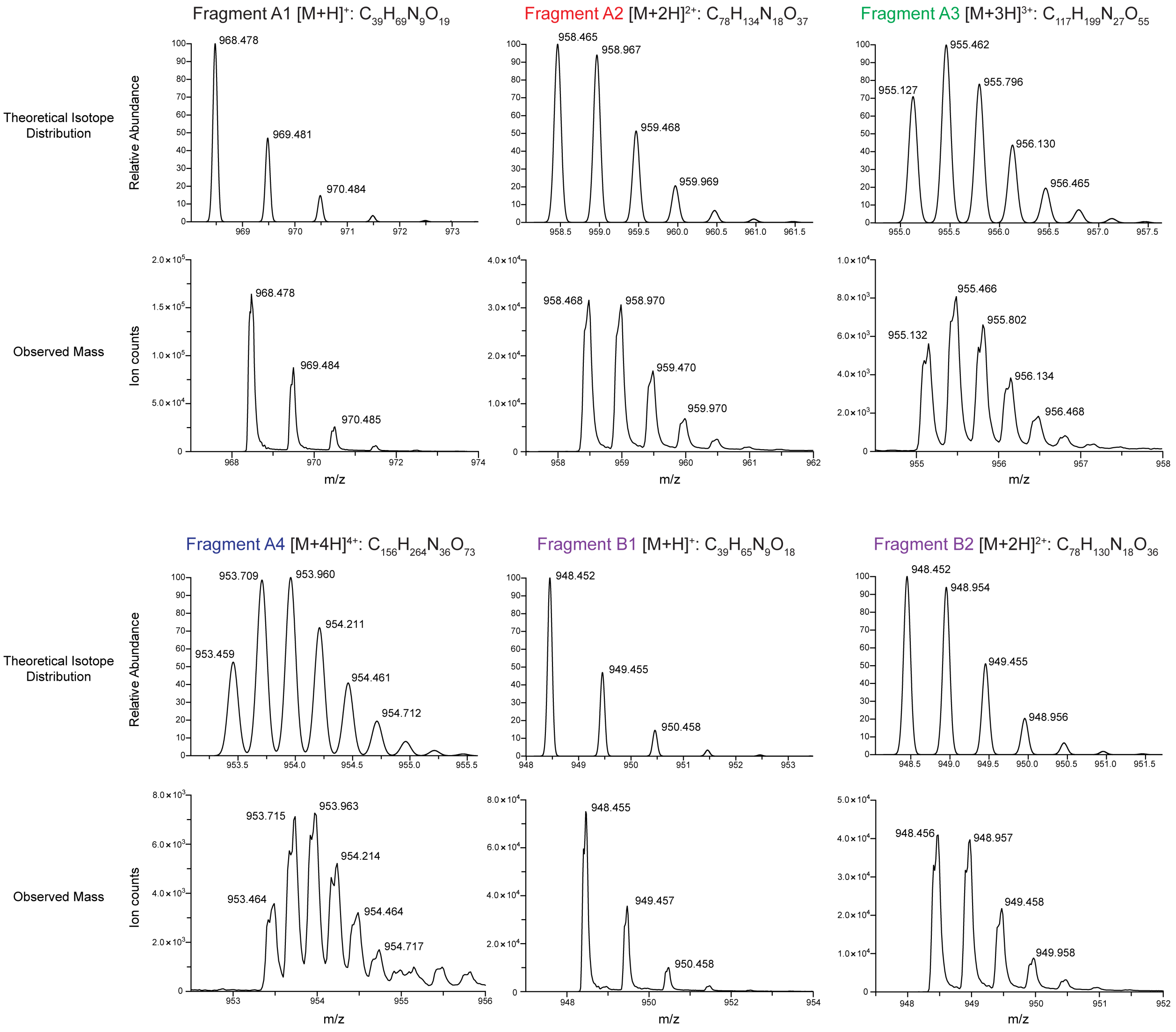


**Supplementary Figure 4. Representative mass spectra of muropeptide products.** See Supplementary Figure 3 for the chemical structure of each product.

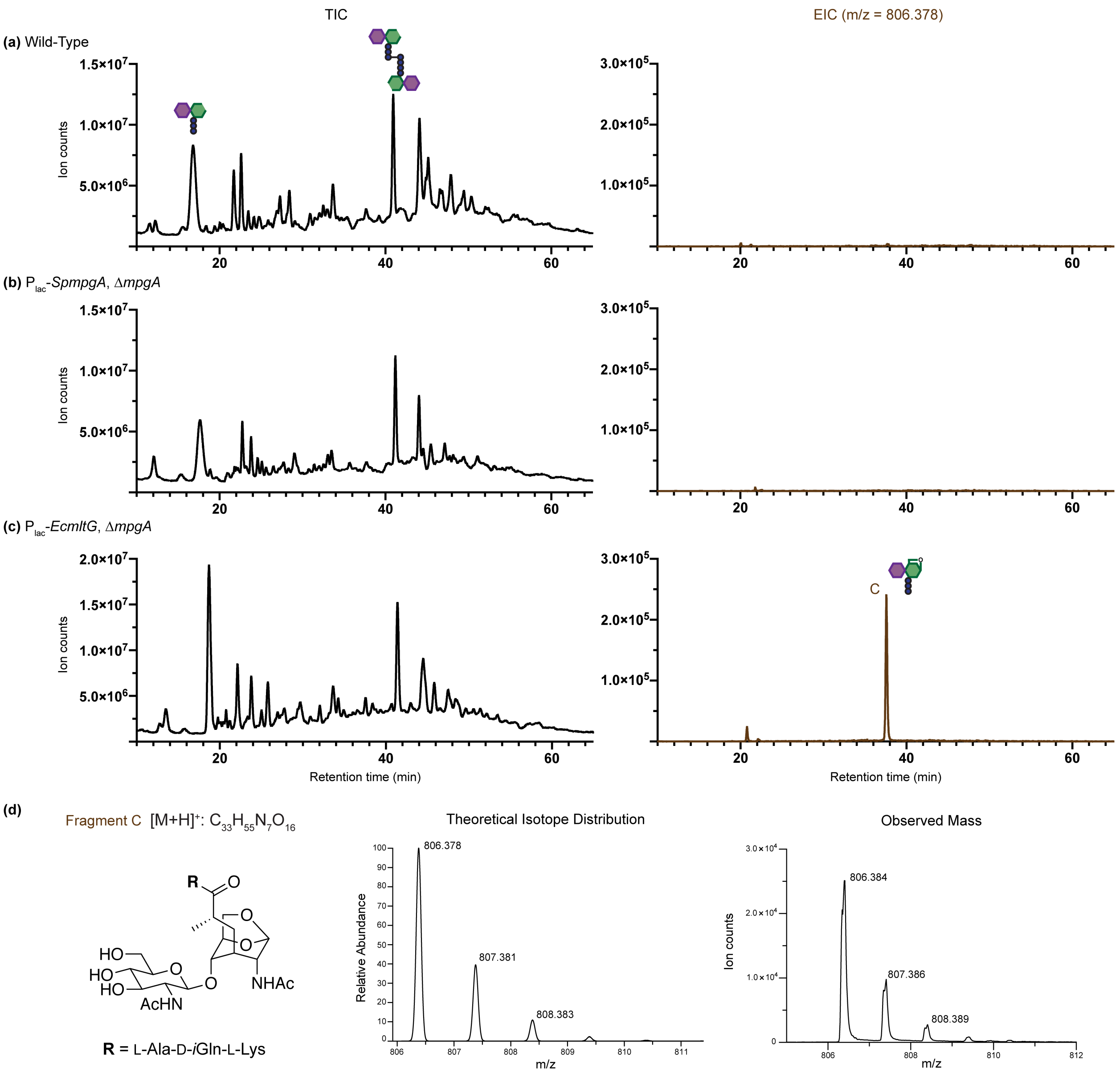


**Supplementary Figure 5. Muropeptides with anhydro-MurNAc ends are detected in peptidoglycan isolated from *S. pneumoniae* complemented with *Ec*MltG, but not in peptidoglycan complemented with *Sp*MpgA.** Isolated peptidoglycan from unencapsulated *S. pneumoniae* D39 (a), $P_{lac}$-*SpmpgA* Δ*mpgA* (b), or $P_{lac}$-*EcmltG* Δ*mpgA* (c) was digested with mutanolysin. The resulting muropeptide products were separated and detected by LC-MS. The total ion chromatogram (TIC) and the extracted ion chromatogram (EIC: m/z = 806.378) are shown for each sample. Major peaks at ~18 min and ~41 min correspond to a tripeptide monomer and crosslinked dimer with unbranched stem peptides, respectively. A peak corresponding to a tripeptide monomer with an anhydro-MurNAc end was only detected in peptidoglycan isolated from cells expressing *Ec*MltG. Note that $P_{lac}$ is a constitutive promoter in the absence of LacI.[1] (d) Chemical structure and mass spectrum of the anhydro-MurNAc containing muropeptide species.

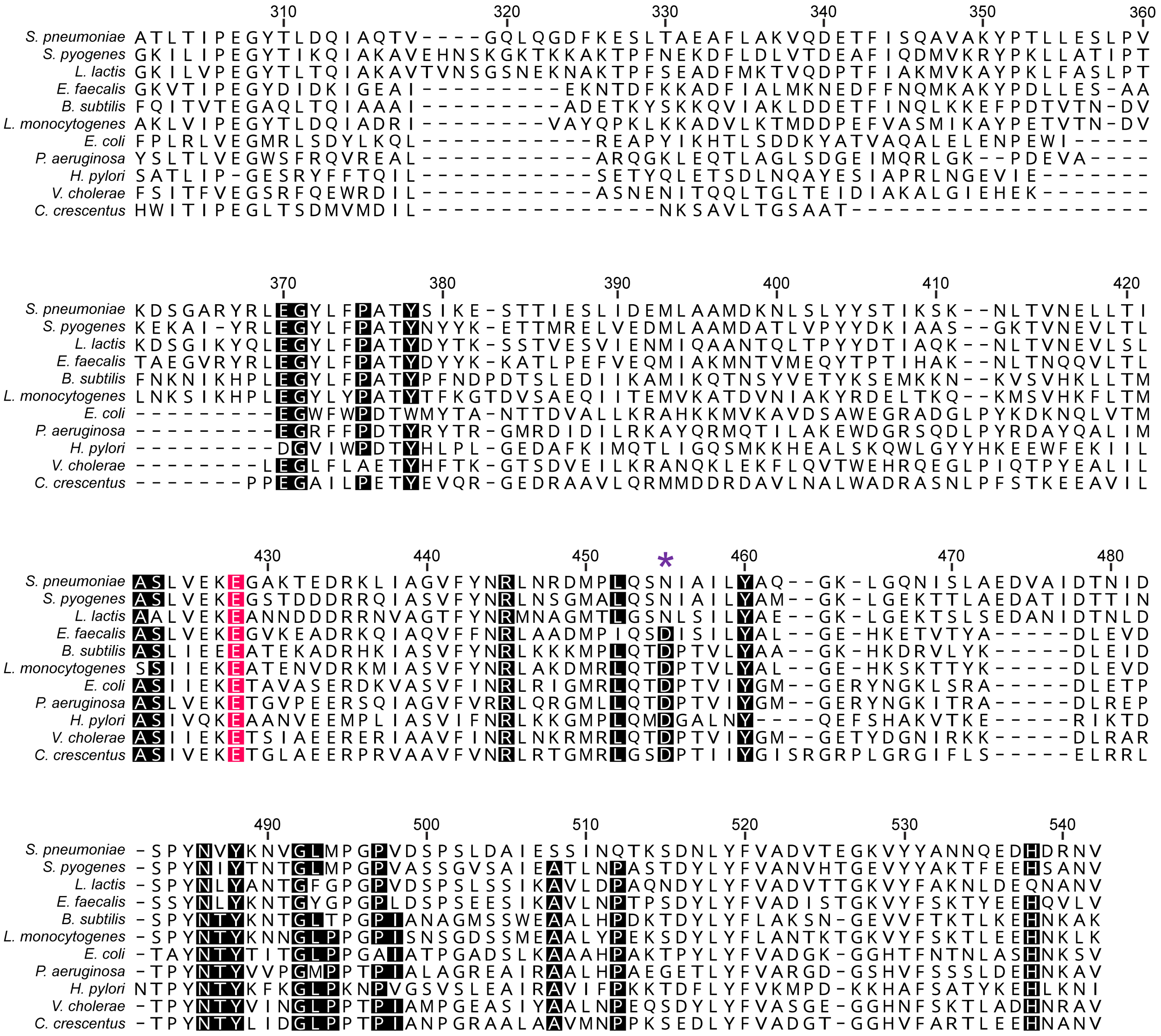


**Supplementary Figure 6. Sequence alignment of the YceG catalytic subdomain.** Sequence conservation analysis of ~15000 sequences containing the YceG domain was performed using the EVcouplings server.[2] Representative examples from 11 species are shown. The catalytic glutamate (~100% conserved) is highlighted in red. Residues conserved in >90% of the analyzed sequences are highlighted in black. The purple asterisk denotes the residue investigated in this study.

**Supplementary Methods**

**Plasmid construction for protein expression**

*Sp*MpgB. The *SPD_0912 (M1-G204)* gene encoding MpgB was amplified from *S. pneumoniae* D39 Δ*cps* genomic DNA using primers oAT101/oAT102. After digestion with NdeI and BamHI, the PCR product was ligated into pET28b. The resulting plasmid pATPL240 expresses $His_6$-*Sp*MpgB. Plasmid pATPL368, expressing $His_6$-*Sp*MpgB$^{D68N}$ (catalytically inactive mutant *Sp*MpgB*) was PCR generated from pATPL240 with oAT103/oAT104 containing the desired mutation.

*St*MpgB. The *stu0757 (A2-L221)* gene encoding MpgB was amplified from *S. thermophilus* LMG18311 genomic DNA using primers oAT105/oAT106, and this fragment was ligated to a linearized pMS211 plasmid backbone (oAT107/oAT108) via InFusion Cloning (TakaraBio). The resulting plasmid pATPL424 expresses $His_6$-SUMO-FLAG-*St*MpgB.

*Ef*MpgB. The *EF1518 (D2-N209)* gene encoding MpgB was amplified from *E. faecalis* V583 genomic DNA using primers oAT109/oAT110 and this fragment was ligated to a linearized pMS211 plasmid backbone (oAT107/oAT108) via InFusion Cloning. The resulting plasmid pATPL425 expresses $His_6$-SUMO-FLAG-*Ef*MpgB.

*Bs*MpgB. The *BSU19130 (K2-E225)* gene encoding MpgB was amplified from *B. subtilis* PY79 genomic DNA using primers oAT111/oAT112 and this fragment was ligated to a linearized pMS211 plasmid backbone (oAT107/oAT108) via InFusion Cloning. The resulting plasmid pATPL426 expresses $His_6$-SUMO-FLAG-*Bs*MpgB.

*Sp*MpgA. The *SPD_1346 (M1-N551)* gene encoding MpgA was amplified from *S. pneumoniae* D39 Δ*cps* genomic DNA using primers oAT113/oAT114. After digestion with NdeI and XhoI, the PCR product was ligated into pET28b. The resulting plasmid pATPL434 expresses $His_6$-*Sp*MpgA. Primer pairs oAT115/oAT116, and oAT117/oAT118 were used to generate expression plasmids pATPL472 ($His_6$-*Sp*MpgA$^{E428Q}$; catalytically inactive mutant *Sp*MpgA*) and pATPL473 ($His_6$-*Sp*MpgA$^{N455D}$), respectively, from pATPL434 via InFusion Cloning. Plasmid pATPL488, expressing $His_6$-*Sp*MpgA$^{\Delta D219\text{-}P295}$ ($His_6$-*Sp*MpgA$^{\Delta LysM}$), was generated from pATPL434 with oAT119/120.

*Ec*MltG. The *b1097 (M1-Q340)* gene encoding MltG was amplified from *E. coli* BL21(DE3) genomic DNA using primers oAT121/oAT122. After digestion with NdeI and XhoI, the PCR product was ligated into pET28b. The resulting plasmid pATPL474 expresses $His_6$-*Ec*MltG. Primer pairs oAT123/oAT124 and oAT125/oAT126 were used to construct expression plasmids pATPL475 (His$_6$-*Ec*MltG$^{E224Q}$; catalytically inactive mutant *Ec*MltG*) and pATPL476 (His$_6$-*Ec*MltG$^{D245N}$), respectively, from pATPL474 via InFusion Cloning.

*Bs*MltG. The *BSU27370 (M1-K360)* gene encoding MltG was amplified from *B. subtilis* PY79 genomic DNA using primers oAT127/oAT128. After digestion with NdeI and XhoI, the PCR product was ligated into pET28b. The resulting plasmid pATPL489 expresses His$_6$-*Bs*MltG.

***S. pneumoniae* strain construction**

AT513 (Δ*lytA*::*kan*): The ~1kb upstream and downstream regions of *lytA* were amplified using primer pairs oAT129/oAT130 and oAT131/oAT132. The *kan* gene was amplified by oAT133/oAT134 from D39 Δ*cps* Δ*bgaA*::*kan* genomic DNA. These PCR fragments were assembled by overlap extension PCR. The resulting PCR cassette was transformed into D39 Δ*cps* and transformants were selected with kanamycin. Integration into the genome was confirmed by diagnostic PCR using primers oAT135 and oAT134.

AT514 (Δ*lytB*::*kan*): The ~1kb upstream and downstream regions of *lytB* were amplified using primer pairs oAT136/oAT137 and oAT138/oAT139. These PCR fragments and *kan* were assembled by overlap extension PCR. The resulting PCR cassette was transformed into D39 Δ*cps* and transformants were selected with kanamycin. Integration into the genome was confirmed by diagnostic PCR using primers oAT140 and oAT134.

AT515 (Δ*lytC*::*kan*): The ~1kb upstream and downstream regions of *lytC* were amplified using primer pairs oAT141/oAT142 and oAT143/oAT144. These PCR fragments and *kan* were assembled by overlap extension PCR. The resulting PCR cassette was transformed into D39 Δ*cps* and transformants were selected with kanamycin. Integration into the genome was confirmed by diagnostic PCR using primers oAT145 and oAT134.

AT516 (Δ*cbpD*::*kan*): The ~1kb upstream and downstream regions of *cbpD* were amplified using primer pairs oAT146/oAT147 and oAT148/oAT149. These PCR fragments and *kan* were assembled by overlap extension PCR. The resulting PCR cassette was transformed into D39 Δ*cps* and transformants were selected with kanamycin. Integration into the genome was confirmed by diagnostic PCR using primers oAT150 and oAT134.

AT520 (Δ*spd_0873*::*erm*): The ~1kb upstream and downstream regions of *spd_0873* were amplified using primer pairs oAT151/oAT152 and oAT153/oAT154. The *erm* gene was amplified by oAT133/oAT134 from D39 Δ*cps* Δ*bgaA*::*erm* genomic DNA. These PCR fragments were assembled by overlap extension PCR. The resulting PCR cassette was transformed into D39 Δ*cps* and transformants were selected with erythromycin. Integration into the genome was confirmed by diagnostic PCR using primers oAT155 and oAT134.

AT446 (Δ*mpgB*): $P_{96}$ (*spd_0104* promoter region) and *B. subtilis sacB* gene were amplified using primer pairs oAT156/oAT157 and oAT158/oAT159 and combined to make a $P_{96}$-*sacB* DNA fragment[3]. The ~1kb upstream and downstream regions of *mpgB* were amplified using primer pairs oAT160/oAT161 and oAT162/oAT163. These PCR fragments and *erm* were assembled by overlap extension PCR. The resulting PCR cassette was transformed into D39 Δ*cps* and transformants were selected with erythromycin (AT428). For making a markerless deletion, the ~1kb upstream and downstream regions of *mpgB* were amplified using primer pairs oAT160/oAT164 and oAT165/oAT163. These PCR fragments were combined by overlap extension PCR, and the resulting PCR cassette was transformed into AT428. Transformants were selected with 10% sucrose, and the loss of resistance marker was assessed by diagnostic PCR using primers oAT166 and oAT167. Markerless deletion of *mpgB* was confirmed by DNA sequencing.

AT563 & AT564: Wild-type (AT563) and Δ*mpgB* (AT564) constitutively expressing LacI were constructed by transforming pPEPY-PF6-lacI to D39 Δ*cps* or AT446 and selecting transformants with gentamycin.

AT580-584: Plasmids containing $P_{lac}$-*mpgB* were constructed by first linearizing the pPEPZ-$P_{lac}$ vector using primers oAT168 and oAT169. This linearized vector was ligated to *SpmpgB* (oAT170/oAT171), *StmpgB* (oAT172/oAT173), *EfmpgB* (oAT174/oAT175) or *BsmpgB* (oAT176/oAT177) via InFusion Cloning to generate pATPL461 ($P_{lac}$-*SpmpgB*), pATPL462 ($P_{lac}$-*StmpgB*), pATPL463 ($P_{lac}$-*EfmpgB*) or pATPL464 ($P_{lac}$-*BsmpgB*). These plasmids were transformed into AT564 and transformants were selected with spectinomycin.

AT600 & AT635: Plasmids containing $P_{lac}$-*EcmltG* or $P_{lac}$-*SpmpgA* were constructed by ligating the linearized pPEPZ-$P_{lac}$ vector to *EcmltG* (oAT178/oAT179) or *SpmpgA* (oAT180/oAT181) via InFusion Cloning. The resulting plasmids pATPL471 ($P_{lac}$-*EcmltG*) and pATPL492 ($P_{lac}$-*SpmpgA*) were transformed into D39 Δ*cps* to generate AT592 and AT634, respectively. A *mpgA* deletion cassette was assembled by first amplifying the ~1kb upstream and downstream regions of *mpgA* using primer pairs oAT182/oAT183 and oAT184/oAT185. These PCR fragments and *kan* were assembled by overlap extension PCR. The resulting PCR cassette was transformed into AT592 or AT634 and transformants were selected with kanamycin. Integration of the deletion cassette into the genome was confirmed by diagnostic PCR using primers oAT186 and oAT134.

**Supplementary Table 1. Bacterial strains used in this study**

| Strain | Description* | Reference |
|---|---|---|
| *E. coli* | | |
| XL1-Blue | Host strain for plasmid cloning | Stratagene |
| Stellar | Host strain for plasmid cloning | TakaraBio |
| C43(DE3) | BL21(DE3) derivative strain for protein production | 4 |
| *S. pneumoniae* | | |
| D39 Δ*cps* | Unencapsulated D39 derivative strain (wild-type) | 5 |
| AT030 | D39 Δ*cps*, Δ*bgaA*::*erm*; Erm$^{R}$ | 6 |
| AT031 | D39 Δ*cps*, Δ*bgaA*::*kan*; Kan$^{R}$ | 6 |
| AT513 | D39 Δ*cps*, Δ*lytA*::*kan*; Kan$^{R}$ | This study |
| AT514 | D39 Δ*cps*, Δ*lytB*::*kan*; Kan$^{R}$ | This study |
| AT515 | D39 Δ*cps*, Δ*lytC*::*kan*; Kan$^{R}$ | This study |
| AT516 | D39 Δ*cps*, Δ*cbpD*::*kan*; Kan$^{R}$ | This study |
| AT520 | D39 Δ*cps*, Δ*spd_0873*::*erm*; Erm$^{R}$ | This study |
| AT446 | D39 Δ*cps*, Δ*mpgB* | This study |
| AT563 | D39 Δ*cps*, Δ*prsA*::(PF6-*lacI*, *gent*); Gent$^{R}$ | This study |
| AT564 | D39 Δ*cps*, Δ*prsA*::(PF6-*lacI*, *gent*), Δ*mpgB*; Gent$^{R}$ | This study |
| AT580 | D39 Δ*cps*, Δ*prsA*::(PF6-*lacI*, *gent*), Δ*mpgB*, Δ*spd_1735*::(P$_{lac}$, *spec*); Gent$^{R}$, Spec$^{R}$ | This study |
| AT581 | D39 Δ*cps*, Δ*prsA*::(PF6-*lacI*, *gent*), Δ*mpgB*, Δ*spd_1735*::(P$_{lac}$-*SpmpgB*, *spec*); Gent$^{R}$, Spec$^{R}$ | This study |
| AT582 | D39 Δ*cps*, Δ*prsA*::(PF6-*lacI*, *gent*), Δ*mpgB*, Δ*spd_1735*::(P$_{lac}$-*StmpgB*, *spec*); Gent$^{R}$, Spec$^{R}$ | This study |
| AT583 | D39 Δ*cps*, Δ*prsA*::(PF6-*lacI*, *gent*), Δ*mpgB*, Δ*spd_1735*::(P$_{lac}$-*EfmpgB*, *spec*); Gent$^{R}$, Spec$^{R}$ | This study |
| AT584 | D39 Δ*cps*, Δ*prsA*::(PF6-*lacI*, *gent*), Δ*mpgB*, Δ*spd_1735*::(P$_{lac}$-*BsmpgB*, *spec*); Gent$^{R}$, Spec$^{R}$ | This study |
| AT600 | D39 Δ*cps*, Δ*mpgA*::*kan,* Δ*spd_1735*::(P$_{lac}$-*SpmpgA*, *spec*); Spec$^{R}$, Kan$^{R}$ | This study |
| AT635 | D39 Δ*cps*, Δ*mpgA*::*kan,* Δ*spd_1735*::(P$_{lac}$-*EcmltG*, *spec*); Spec$^{R}$, Kan$^{R}$ | This study |

*Abbreviations: Amp$^{R}$, ampicillin/carbenicillin resistance; Cm$^{R}$, chloramphenicol resistance; Erm$^{R}$, erythromycin resistance; Gent$^{R}$, gentamycin resistance; Kan$^{R}$, kanamycin resistance; Spec$^{R}$, spectinomycin resistance

**Supplementary Table 2. Plasmids used in this study**

| Plasmid | Description* | Reference |
|---|---|---|
| pET28b(+) | IPTG-inducible protein expression vector; $Kan^R$ | Novagen |
| pATPL240 | $His_6$-*Sp*MpgB expression vector; $Kan^R$ | This study |
| pATPL368 | $His_6$-*Sp*MpgB$^{D68N}$ expression vector; $Kan^R$ | This study |
| pATPL434 | $His_6$-*Sp*MpgA expression vector; $Kan^R$ | This study |
| pATPL472 | $His_6$-*Sp*MpgA$^{E428Q}$ expression vector; $Kan^R$ | This study |
| pATPL473 | $His_6$-*Sp*MpgA$^{N455D}$ expression vector; $Kan^R$ | This study |
| pATPL488 | $His_6$-*Sp*MpgA$^{\Delta D219\text{-}P295}$ expression vector; $Kan^R$ | This study |
| pATPL474 | $His_6$-*Ec*MltG expression vector; $Kan^R$ | This study |
| pATPL475 | $His_6$-*Ec*MltG$^{E224Q}$ expression vector; $Kan^R$ | This study |
| pATPL476 | $His_6$-*Ec*MltG$^{D245N}$ expression vector; $Kan^R$ | This study |
| pATPL489 | $His_6$-*Bs*MltG expression vector; $Kan^R$ | This study |
| pMS211 | $His_6$-SUMO-FLAG-*Tt*RodA expression vector; $Amp^R$ | 7 |
| pATPL424 | $His_6$-SUMO-FLAG-*St*MpgB expression vector; $Amp^R$ | This study |
| pATPL425 | $His_6$-SUMO-FLAG-*Ef*MpgB expression vector; $Amp^R$ | This study |
| pATPL426 | $His_6$-SUMO-FLAG-*Bs*MpgB expression vector; $Amp^R$ | This study |
| pAM174 | Encodes arabinose-inducible Ulp1$^{L403\text{-}K621}$ protease; $Cm^R$ | 8 |
| pMgt1 | *S. aureus* SgtB-$His_6$ expression vector; $Amp^R$ | 9 |
| pET24bSgtBY181D | *S. aureus* SgtB$^{Y181D}$-$His_6$ expression vector; $Kan^R$ | 10 |
| pPEPY-PF6-lacI | *S. pneumoniae prsA::*PF6-*lacI* integration vector; $Gent^R$, $Kan^R$ | 11, Addgene |
| pPEPZ-$P_{lac}$ | *S. pneumoniae spd_1735* integration vector containing IPTG-inducible $P_{lac}$ promoter; $Spec^R$ | 1, Addgene |
| pATPL461 | *S. pneumoniae spd_1735*::$P_{lac}$-*SpmpgB* integration vector; $Spec^R$ | This study |
| pATPL462 | *S. pneumoniae spd_1735*::$P_{lac}$-*StmpgB* integration vector; $Spec^R$ | This study |
| pATPL463 | *S. pneumoniae spd_1735*::$P_{lac}$-*EfmpgB* integration vector; $Spec^R$ | This study |
| pATPL464 | *S. pneumoniae spd_1735*::$P_{lac}$-*BsmpgB* integration vector; $Spec^R$ | This study |
| pATPL471 | *S. pneumoniae spd_1735*::$P_{lac}$-*EcmltG* integration vector; $Spec^R$ | This study |
| pATPL492 | *S. pneumoniae spd_1735*::$P_{lac}$-*SpmpgA* integration vector; $Spec^R$ | This study |

*Abbreviations: $Amp^R$, ampicillin/carbenicillin resistance; $Cm^R$, chloramphenicol resistance; $Erm^R$, erythromycin resistance; $Gent^R$, gentamycin resistance; $Kan^R$, kanamycin resistance; $Spec^R$, spectinomycin resistance

**Supplementary Table 3. Oligonucleotide primers used in this study**

| Primer | Sequence (5'-3')* |
|---|---|
| oAT101 | GTACATATGTTTAAACGAATTCGAAGAGTGCTTGT |
| oAT102 | ACTGGATCCCTAGCCAGATGTTGAAAA |
| oAT103 | AATGTTATGCAGTCTAGTGAGTCT |
| oAT104 | GCCTTCTTTTCCTTTTGTTTCAGTATA |
| oAT105 | CCTGGGGGGTCATCCTTTAAATGGATAAGACGTCTGGTGGT |
| oAT106 | TGCAGTCACCCGGGCTTAGAGACGAGAGAATATTCGTATCAGAAAG |
| oAT107 | GCCCGGGTGACTGCAGGA |
| oAT108 | GGATGACCCCCCAGGGCC |
| oAT109 | CCTGGGGGGTCATCCGATGATTCGATGCGTAGAGTAAGA |
| oAT110 | TGCAGTCACCCGGGCTTAATTTAACTTTTCAATAAACCAACGATTC |
| oAT111 | CCTGGGGGGTCATCCAAGAAAAAGAGAAAAGGCTGTTTC |
| oAT112 | TGCAGTCACCCGGGCTTATTCATGAGCCTTGGATTCC |
| oAT113 | GTACATATGAGTGAAAAGTCAAGAGAAGAAGAGA |
| oAT114 | ACTCCTCGAGTTAGTTTAATTTGCTGTTGACATGTTCAG |
| oAT115 | TCGAAAAACAAGGTGCCAAGACAGAAGATCG |
| oAT116 | CACCTTGTTTTTCGACCAAGGAAGCAATGG |
| oAT117 | TTCAAAGTGATATTGCAATCTTGTATGCCCAAGG |
| oAT118 | CAATATCACTTTGAAGTGGCATATCACG |
| oAT119 | GATAGGTAATAAGGAATCTAGCACGTACT |
| oAT120 | CAAGAACCTGTACTTGCGACTTTG |
| oAT121 | AGTACATATGAAAAAAGTGTTATTGATAATCTTGTTATT |
| oAT122 | ACTCCTCGAGTTACTGCGCATTTTTTTCCTTAAG |
| oAT123 | TCGAAAAACAAACCGCCGTTGCCAGTG |
| oAT124 | CGGTTTGTTTTTCGATAATTGATGCCATCGTC |
| oAT125 | TGCAGACCAACCCGACCGTGATTTACGGG |
| oAT126 | TCGGGTTGGTCTGCAGGCGCATACC |
| oAT127 | AGTACATATGTATATCAATCAGCAAAAAAAATCGTTT |
| oAT128 | ACTCCTCGAGTTATTTCTCATTTTTTGAGGAAATGTATTTTTC |
| oAT129 | ATGAGTTCAATTGTATCTATCGGCAG |
| oAT130 | TCCTGCCTTTCCTCCCTCATTCTACTCCTTATCAATTAAAACAACTC |
| oAT131 | GAGAGCACAGATACGGCGTAATGGAATGTCTTTCAAATCAGAACAG |
| oAT132 | TCCTCAATCTATATAACATAGCTTTATGAC |
| oAT133 | GAGGGAGGAAAGGCAGGA |
| oAT134 | CGCCGTATCTGTGCTCTC |
| oAT135 | CGGTTCTCTGCTTTTATTATATTCG |
| oAT136 | CACAGGAACAGTTGTATTATAAGGAG |
| oAT137 | TCCTGCCTTTCCTCCCTCATTACAAACTAATAAAAAATCAAGAACAGAT |
| oAT138 | GAGAGCACAGATACGGCGTACTATAAGTGAATATGATTTGAGTGAATAG |
| oAT139 | ATGCTTGACTGAGACTTCCTTCA |

* Underlined sequences are restriction sites introduced in primers.

**Supplementary Table 3. Oligonucleotide primers used in this study (cont.)**

| Primer | Sequence (5'-3') |
|---|---|
| oAT140 | TCTCTGGAATCTCGTATGGC |
| oAT141 | AGAAAGAGAAGCAAGCAATCCTCA |
| oAT142 | TCCTGCCTTTCCTCCCTCCTGAATGCTGTTCCACCTAG |
| oAT143 | GAGAGCACAGATACGGCGGCGATGATTTGAAAGAGGGATGT |
| oAT144 | ATTTCTTACAAACCAGGTGCTTG |
| oAT145 | GATGGAACCAGTGCTGACTTGA |
| oAT146 | CTCGAAATGGGTGCGGAAAG |
| oAT147 | TCCTGCCTTTCCTCCCTCTCTTCCTCCTTAAAAAATAATATAAAGCGAT |
| oAT148 | GAGAGCACAGATACGGCGAAAATTGGAGTAGGAGAAATTTCC |
| oAT149 | CATAAAAGTAAGGCAGGCTAACC |
| oAT150 | GAATATCGCTCAGAATATTGGGA |
| oAT151 | CCTTTCAAGATAACATTGGCTTC |
| oAT152 | TCCTGCCTTTCCTCCCTCATCTTATAATTCTACCCTAAAAATCAAAAAAAAT |
| oAT153 | GAGAGCACAGATACGGCGCATTTGTTTATCATTTGCTTTTTCTTTTTG |
| oAT154 | TGATGAGGGATAGCTGCTTTAG |
| oAT155 | CCTTTCACCTCAACAAAGTTACC |
| oAT156 | TCCGATGATATCAAAGACAGATTGAAA |
| oAT157 | ATTCGAAAATTCTCCTTCTTTCTATAGTT |
| oAT158 | ATAGAAAGAAGGAGAATTTTCGAATATGAACATCAAAAAGTTTGCAAAAC |
| oAT159 | TCCTGCCTTTCCTCCCTCTTATTTGTTAACTGTTAATTGTCCTTG |
| oAT160 | ATTCAAGACAAGTGGGATTCGT |
| oAT161 | TTTCAATCTGTCTTTGATATCATCGGATTATTTACTTTGGATATCCTCGATATTTTTGA |
| oAT162 | GAGAGCACAGATACGGCGACCAGGTGTTTTTGTTATAAGTTTTCT |
| oAT163 | CGTTACGGTTACCATCCATTATACC |
| oAT164 | TTATTTACTTTGGATATCCTCGATATTTTTGA |
| oAT165 | TCAAAAATATCGAGGATATCCAAAGTAAATAACCAGGTGTTTTTGTTATAAGTTTTCT |
| oAT166 | TTGCTGGATGAAGTCAATATTACCC |
| oAT167 | CAACTTCTACAATATCATTTTTCTTTAAC |
| oAT168 | TATTTTTCCTCCTTATTTATTTAGATCTTAATTGTG |
| oAT169 | GATCCCTCCAGTAACTCGAG |
| oAT170 | TAAGGAGGAAAAATAATGTTTAAACGAATTCGAAGAGTGCTT |
| oAT171 | GTTACTGGAGGGATCCTAGCCAGATGTTGAAAAGAGAGTG |
| oAT172 | TAAGGAGGAAAAATAATGTTTAAATGGATAAGACGTCTGGTG |
| oAT173 | GTTACTGGAGGGATCTTAGAGACGAGAGAATATTCGTATCAGAAA |
| oAT174 | TAAGGAGGAAAAATAATGGATGATTCGATGCGTAGAGTAA |
| oAT175 | GTTACTGGAGGGATCTTAATTTAACTTTTCAATAAACCAACGATT |
| oAT176 | TAAGGAGGAAAAATAATGAAGAAAAAGAGAAAAGGCTGTTTC |
| oAT177 | GTTACTGGAGGGATCTTATTCATGAGCCTTGGATTCC |
| oAT178 | TAAGGAGGAAAAATAATGAAAAAAGTGTTATTGATAATCTTGTT |

**Supplementary Table 3. Oligonucleotide primers used in this study (cont.)**

| Primer | Sequence (5'-3') |
|---|---|
| oAT179 | GTTACTGGAGGGATCTTACTGCGCATTTTTTTCCTTAAG |
| oAT180 | TAAGGAGGAAAAATAATGAGTGAAAAGTCAAGAGAAGAAGAGA |
| oAT181 | GTTACTGGAGGGATCTTAGTTTAATTTGCTGTTGACATGTTCAG |
| oAT182 | ATCATTCAGGCAAGCAAGTCT |
| oAT183 | TCCTGCCTTTCCTCCCTCAAGTTTTTCCTCCTTGTTGATAATCC |
| oAT184 | GAGAGCACAGATACGGCGTAAACAAACTAAAATTATGTGATACTTCA |
| oAT185 | AATCTAAAGTATAGTGAAATGAAATAAAACATG |
| oAT186 | TTTACGTCTTTATGGGAGCAG |

### References


1. Keller, L. E., Rueff, A.-S., Kurushima, J. & Veening, J.-W. Three New Integration Vectors and Fluorescent Proteins for Use in the Opportunistic Human Pathogen Streptococcus pneumoniae. *Genes (Basel).* **10**, 394 (2019).

2. Hopf, T. A. *et al.* The EVcouplings Python framework for coevolutionary sequence analysis. *Bioinformatics* **35**, 1582–1584 (2019).

3. Lo Sapio, M., Hilleringmann, M., Barocchi, M. A. & Moschioni, M. A novel strategy to over-express and purify homologous proteins from Streptococcus pneumoniae. *J. Biotechnol.* **157**, 279–286 (2012).

4. Miroux, B. & Walker, J. E. Over-production of proteins in Escherichia coli: Mutant hosts that allow synthesis of some membrane proteins and globular proteins at high levels. *J. Mol. Biol.* **260**, 289–298 (1996).

5. Lanie, J. A. *et al.* Genome sequence of Avery's virulent serotype 2 strain D39 of Streptococcus pneumoniae and comparison with that of unencapsulated laboratory strain R6. *J. Bacteriol.* **189**, 38–51 (2007).

6. Fenton, A. K., Mortaji, L. El, Lau, D. T. C., Rudner, D. Z. & Bernhardt, T. G. CozE is a member of the MreCD complex that directs cell elongation in Streptococcus pneumoniae. *Nat. Microbiol.* **2**, 16237 (2016).

7. Sjodt, M. *et al.* Structure of the peptidoglycan polymerase RodA resolved by evolutionary coupling analysis. *Nature* **556**, 118–121 (2018).

8. Meeske, A. J. *et al.* SEDS proteins are a widespread family of bacterial cell wall polymerases. *Nature* **537**, 634–638 (2016).

9. Heaslet, H., Shaw, B., Mistry, A. & Miller, A. A. Characterization of the active site of S. aureus monofunctional glycosyltransferase (Mtg) by site-directed mutation and structural analysis of the protein complexed with moenomycin. *J. Struct. Biol.* **167**, 129–135 (2009).

10. Rebets, Y. *et al.* Moenomycin resistance mutations in Staphylococcus aureus reduce peptidoglycan chain length and cause aberrant cell division. *ACS Chem. Biol.* **9**, 459–467 (2014).

11. Liu, X. *et al.* High-throughput CRISPRi phenotyping identifies new essential genes in Streptococcus pneumoniae. *Mol. Syst. Biol.* **13**, 931 (2017).
